# Electrical stimulation combined with p27Kip1 inactivation drives proliferative neurogenic reprogramming of Mueller glia in the adult mouse retina

**DOI:** 10.64898/2026.02.05.704102

**Authors:** Megan L. Stone, Joel Jovanovic, Edward M. Levine

**Author notes:** **Corresponding author:** Edward M. Levine, B3222 Medical Center North 1161 21st Ave South Nashville, Tennessee 37232.

## Abstract

Mueller glial reprogramming studies demonstrate that mammalian Mueller glia can be induced to proliferate and/or engage in neural differentiation, as occurs naturally in teleost fish. A major objective is the identification of combined strategies that promote both robust proliferation and neurogenesis. These studies would benefit from a translatable screening platform that enables controlled perturbation, maintained tissue context and longitudinal analysis, such as 3D culture for first tier analysis of reprogramming strategies. Here, we validate a 3D retinal culture for Mueller glial reprogramming studies by recapitulating key signatures of an *in vivo* reprogramming paradigm. Next, we find that electrical stimulation (E-stim) as a tunable, extrinsic cue is sufficient to activate endogenous Ascl1 expression, indicating a state transition favorable for neurogenesis, while Mueller glia-specific *p27^Kip^*^1^ inactivation promotes robust, prolonged proliferation. Utilization of the lineage-tracing proliferation-history reporter H3.1-iCOUNT enabled longitudinal proliferation analysis and assessment of reprogramming outcomes within the proliferative, Mueller-derived population. With this model, we find that E-stim and *p27^Kip^*^1^ inactivation in combination (ESPI) increases proliferation, endogenous Ascl1 expression, and neurogenesis of Mueller-derived cells across modalities. Together, this work establishes a 3D culture framework for discovery of combinatorial reprogramming strategies within a proliferative context and identifies ESPI as an efficient approach to proliferative, neurogenic Mueller glial reprogramming.

## Introduction

The adult mammalian retina lacks an endogenous, replicative progenitor population capable of replacing neurons after injury. Mueller glia are a promising candidate for autologous cell replacement, motivated by robust regenerative programs in fish. Initially discovered in goldfish and extensively studied in zebrafish, Mueller glia reliably exit quiescence, proliferate, and generate multipotent progenitors that replenish neurons to restore tissue function after injury^1–3^. By contrast, mammalian Mueller glia adopt a reactive state with limited proliferative responses and rare, if any, neurogenic conversion, indicating constraints to neural regeneration^4^.

A central challenge in mammalian Mueller glia-mediated regeneration is identifying conditions that sequentially drive productive proliferation and acquisition of neurogenic competence. Robust proliferation is an important piece of the zebrafish regeneration paradigm, as it is required to produce enough cells to restore tissue function without depleting the Mueller glia population, and may open a window of plasticity that could be permissive to neurogenic conversion with additional cues^5^. Combinatorial strategies have emerged that integrate validated approaches to induce proliferation with drivers of neurogenic differentiation, including YAP overexpression with NFI factor inactivation^6^, and inhibited monocyte invasion with ASCL1 overexpression^7^. These studies demonstrate that combined strategies to promote proliferation and neurogenic programs can be engaged concurrently in mammalian Mueller glia to improve efficiency of these outcomes.

As combinatorial approaches to Mueller glial reprogramming become more complex, models that preserve relevant biological context while enabling efficient evaluation of reprogramming outcomes are needed to alleviate the burden of the time, technical challenges, and animal requirements that scales with iterative testing *in vivo*^8^. To this end, 3D culture of adult retina tissue offers streamlined analysis, facilitates longitudinal tracking of proliferation, morphology, and state change marker expression, and is a direct model to optimize conditions for translation to the human *ex vivo* retina^9,10^. While explant viability can limit testing duration, Mueller glia remain viable and tractable through the experimental timelines used here. Notably, current 3D culture mimics an injury-like context relevant to regenerative reprogramming^8,11–13^.

In this study, we validate 3D adult mouse retina culture as a model for Mueller glial reprogramming through recapitulation of the combinatorial ASCL1 overexpression and small molecule reprogramming paradigm ANTSi elucidated *in vivo*^14^. We then report a new reprogramming paradigm that combines electrical stimulation (E-stim) as an extrinsic cue to drive neurogenic reprogramming and genetic inactivation of the cyclin-dependent kinase inhibitor gene *p27^Kip^*^1^ (gene symbol: *Cdkn1b*; referred to as p27) to induce cell-cycle release for Mueller glia proliferation. Separately, E-stim was sufficient to induce endogenous expression of Ascl1 and Otx2, demonstrating a fate change favorable for neurogenesis, while p27 inactivation efficiently induced prolonged proliferation with indications of neurogenic competence in some cells. Incorporation of a lineage-tracing, proliferation history tracking model allowed for longitudinal proliferation assessment and downstream analysis within the enriched, proliferative, Mueller-derived population. Combined, E-stim and p27 inactivation promoted activation of neurogenic reprogramming associated with induction of endogenous Ascl1 expression in proliferative mammalian Mueller glia with higher efficiency than either approach alone. This study defines an accessible platform for testing existing reprogramming paradigms and discovering new approaches to activating neurogenic programs in proliferative Mueller glia.

## Results

### A modified ANTSi protocol in 3D adult retinal culture promotes Mueller glia reprogramming

The ANTSi paradigm combines *Ascl1* overexpression, HDAC inhibition via Trichostatin-A (TSA), and Stat inhibition via SH-4-54 in the NMDA-injured retina to reprogram Mueller glia into bipolar-like interneurons^14,15^. Key markers of reprogramming included the homeodomain transcription factor Otx2 and calcium binding protein Cabp5 as evidence of neuronal conversion and bipolar differentiation. Otx2 is expressed in bipolar cells and photoreceptors and is required for their fate specification and differentiation during development^16^, and Cabp5 is expressed in rod- and subsets of cone-bipolar cells and modulates synaptic transmission^17^. A low level of BrdU incorporation was also observed in ANTSi, indicating cell cycle reentry but with limited evidence of productive proliferation^7^. Early studies tested *Ascl1* overexpression in a 3D retinal model from very young mice (P12-P14)^18,19^, after which ANTSi was elucidated for the adult retina *in vivo*^14,15^. To determine if 3D culture could be used to study reprogramming in adult retinal tissue, ANTSi was adapted here for adult retina cultured in the air-liquid interface culture format (ANTSi *ex vivo*).

Retinal tissue from Mueller glia-specific *Ascl1* overexpressing mice aged 8-12 weeks were placed into culture 7 days after tamoxifen treatment. Since neuronal cell loss occurs rapidly in culture^8^, NMDA was not used to induce degeneration. BrdU, TSA, and SH-4-54 were added to the media with similar timing as done *in vivo* (Figure 1A), and expression of OTX2 and CABP5 were used as readouts of reprogramming after 16 days *ex vivo* (DEV). As bipolar cells and, to a lesser extent, photoreceptors persist in 3D culture^8^, a lineage reporter to track Mueller glia and their progeny is essential. *eGFP* is expressed from the *Ascl1* transgene (Ascl1-ires-eGFP) and from a second recombination reporter (CAG-CAT-eGFP) to boost signal (Figure 1B)^14,15^. For untreated controls, we used our validated reporter line (*Rlbp1-CreER; Rosa^Ai^*^14^; referred to as MG-Tom), which expresses tdTomato in recombined Mueller glia and their progeny (Figure 1C)^20^. The proportion of reporter^+^ cells (Mueller glia or their progeny) that expressed OTX2 was significantly increased in ANTSi *ex vivo-*treated retinas compared to controls (Figure 1D(ii), E), as were the proportions of Mueller glia co-expressing OTX2 and BrdU (Figure 1D(iii), F), or CABP5 and OTX2 (Figure 1D(iv), G). These observations are consistent with key outcomes of ANTSi *in vivo*, confirming that the *ex vivo* retina can be used to evaluate strategies to drive Mueller glia reprogramming.

**Figure 1.**
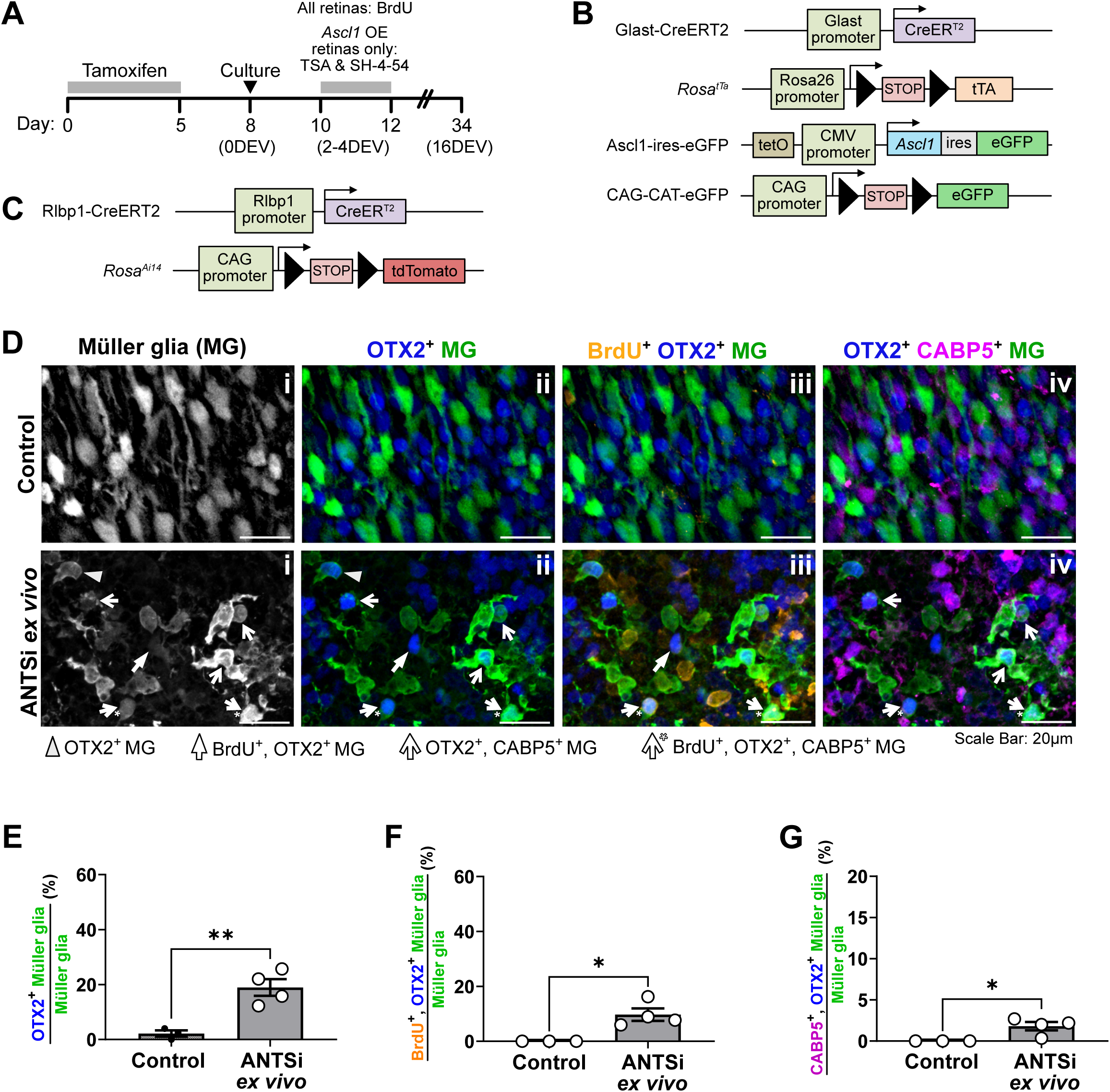
ASCL1-based reprogramming model, ANTSi, recapitulates key reprogramming signatures in the *ex vivo* retina. **(A)** Experimental timeline including tamoxifen induction period *in vivo*, culture establishment (serves as damage model), and drug delivery (BrdU to all conditions; TSA and SH-4-54 to ANTSi *ex vivo* condition only). Tissues fixed at 16 DEV. **(B)** Schematic of ANTSi *ex vivo* lineage reporter model: eGFP-linked ASCL1 expression and additional eGFP expression “boost” are activated in Mueller glia with the Glast:CreER^T2^ driver. Ascl1-ires-eGFP is activated by cre-dependent rTA expression using the TET-off system. **(C)** Control lineage reporter model: TdTomato expression (*Rosa^Ai^*^14^) is activated in Mueller glia with the Rlbp1:CreER^T2^ driver (MG-Tom in text). **(D)** Representative 63x images of whole mount retina showing lineage-traced Mueller glia (i; MG), expression of OTX2 (ii), BrdU incorporation (iii), and CABP5 (iv) in control versus ANTSi *ex vivo* cultures. **(E–G)** Quantification of the proportion of lineage-traced Mueller glia that are OTX2^+^ **(E)**, BrdU^+^ and OTX2^+^ **(F),** or OTX2^+^ and CABP5^+^ **(G)** in control vs. ANTSi *ex vivo* conditions. N=3-4.

### Electrical stimulation promotes cell cycle reentry and limited evidence of reprogramming in Mueller glia

We recently reported that applying five 50 ms square wave electrical pulses (0.042 kV/cm) to adult retinal tissue in air-liquid interface culture promoted proliferation in Mueller glia^13^. To determine if this could induce proliferative reprogramming, MG-Tom retinas (Figure 2A) were stimulated at 14DEV, coinciding with the timing that induced proliferation in our previous study. BrdU was added to the cultures 24 hours after E-stim to label proliferating cells and preclude transient cell cycle reentry that could have resulted from acute injury (Figure 2B). At 19DEV, the proportions of BrdU^+^ (Figure 2C(ii), D) and OTX2^+^ (Figure 2C(iii), E) cells were increased in MG-Tom^+^ cells, suggestive of proliferative reprogramming, but co-labeling with BrdU and OTX2 was rare (Figure 2C(iv), F), and co-labeling with CABP5 or CRX, the latter a homeodomain protein expressed in photoreceptors and bipolar cells and required for their development^21^, was not observed (Figure S1). Thus, if reprogramming occurred, it is unlikely to have been coupled to proliferation or to have progressed to neuronal differentiation, as observed with ANTSi treatment. However, since Otx2 expression in MG-Tom^+^ cells was observed, we asked if endogenous Ascl1 expression was induced following E-stim. Indeed, the number of ASCL1^+^ MG-Tom cells was increased compared to unstimulated controls (Figure 2G-I). Taken together, these observations suggest that E-stim promotes cell cycle re-entry, activation of an endogenous neurogenic reprogramming factor (Ascl1), and expression of a downstream differentiation factor (Otx2) in adult Mueller glia, though not necessarily in a sequential or coordinated manner.

**Figure 2.**
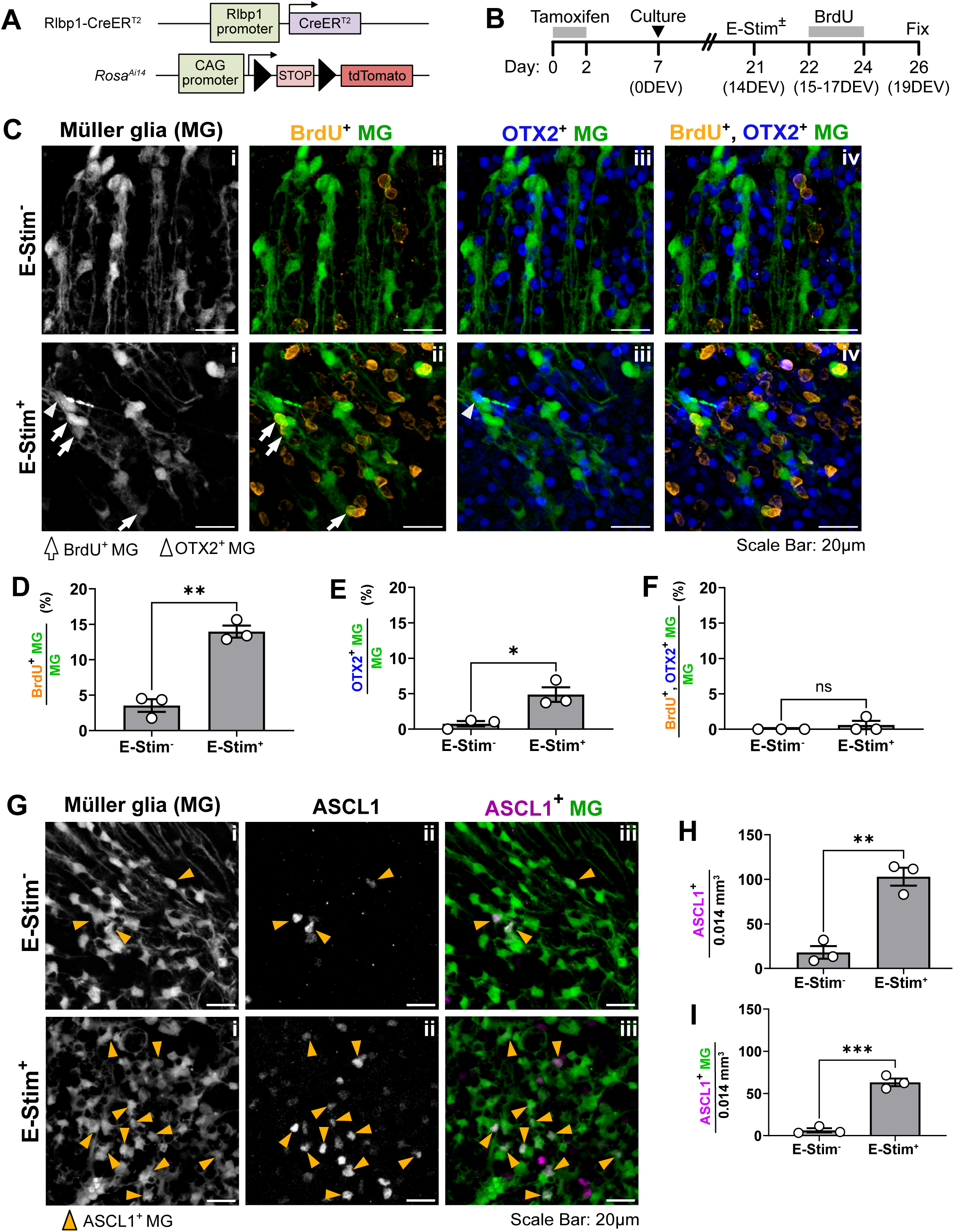
Electrical stimulation promotes uncoupled Mueller glial proliferation and neurogenic reprogramming in Mueller glia. **(A)** Mueller glia-specific lineage reporter model. **(B)** Experimental timeline including reporter induction via tamoxifen, culture establishment, timepoint of electrical stimulation (E-stim^+^) or no treatment in unstimulated control (E-stim^−^), BrdU administration following an excluded 24-hour initial injury response period, and fixation. **(C)** Representative 63x images showing lineage-traced Mueller glia (MG), BrdU incorporation, and OTX2 expression in paired unstimulated (E-stim^−^) and stimulated (E-stim^+^) retinas. **(D–F)** Quantification of Mueller glia co-labeled with BrdU **(D)**, OTX2 **(E)**, or both **(F)** in E-stim^−^ vs. E-stim^+^ conditions. N=3. **(G)** Representative images of ASCL1 immunostaining in lineage-traced Mueller glia in E-stim^−^ vs. E-stim^+^ conditions. **(H,I)** Quantification of ASCL1^+^ cells per ROI volume **(H)** and ASCL1^+^ lineage-traced Mueller glia per ROI **(I)** in paired E-stim^−^ vs. E-stim^+^ retinas. N=3. Statistics in C-E and G-H analyzed using Student’s T-test.

We also observed ASCL1 expression in the unstimulated retina and in MG-Tom-negative cells in both conditions (Figure 2G-I), suggesting that the culture paradigm itself could influence neurogenic competence, albeit at very low efficiency. The latter observation could also be due to Ascl1 activation in Mueller glia that did not express the MG-Tom reporter or because Ascl1 was activated in other, unmarked retinal cell types. Importantly, given the specificity of the MG-Tom reporter system to Mueller glia and their progeny^20^, the localization of Ascl1 and other markers in MG-Tom^+^ cells is strong evidence for changes associated with reprogramming in these cells.

### Mueller glial-specific *p27Kip1* (p27) inactivation combined with E-stim enhanced proliferation and Otx2 expression compared to the individual treatments

p27 is a cyclin-dependent kinase inhibitor protein with key roles in regulating cell cycle progression and maintaining quiescence in proliferation-competent cells. During retinal development p27 coordinates the cell cycle exit of retinal progenitor cells as they initiate differentiation into most retinal cell types including Muller glia^22–25^. p27 is expressed in mature Mueller glia and is transiently downregulated in response to damage coincident with a brief period of cell cycle reentry^26–29^. Conditional genetic inactivation in the adult retina caused Mueller glia to reenter the cell cycle, transiently proliferate, migrate into the photoreceptor layer, and become reactive in the absence of injury^30^. AAV-based Mueller glia-specific *p27* knockdown combined with Cyclin D1 overexpression had similar effects but with rare populations (∼1%) of Mueller glia-lineage labeled cells positive for OTX2 or HuC/D, the latter detecting ELAV RNA binding proteins expressed in amacrine cells^31^. In addition, *p27* conditional genetic inactivation combined with the small molecule LATS inhibitor LKI enhanced Mueller glia proliferation with evidence of rare, though not quantified, Mueller glia lineage-labeled cells positive for Crx^32^. Taken together, these studies support a model in which p27 actively maintains the quiescent or resting Mueller glial state, and its downregulation enables both entry into the cell cycle and a window of cell state plasticity. Whereas the dominant outcome of p27 suppression is glial reactivity in mammalian Mueller glia, neurogenic conversion could be possible with additional intervention. Given our observation that E-stim stimulates proliferation and certain elements of reprogramming, we hypothesized that the combination of E-stim and p27 inactivation could synergize to promote proliferative reprogramming.

To test this, we generated MG-Tom mice containing the *p27^flox^*allele for Mueller glia-specific p27 inactivation and lineage-tracking (Figure 3A). A high tamoxifen dose was applied to maximize recombination and p27 inactivation, and the treatment timeline was similar to E-stim alone (Figure 2B) but with a 48 hr BrdU labeling interval beginning 2 days after E-stim (Figure 3B). The individual and combined effects of E-stim and p27 inactivation were assessed, and untreated MG-Tom retina served as the control (referred to as p27 WT in Figure 3).

**Figure 3.**
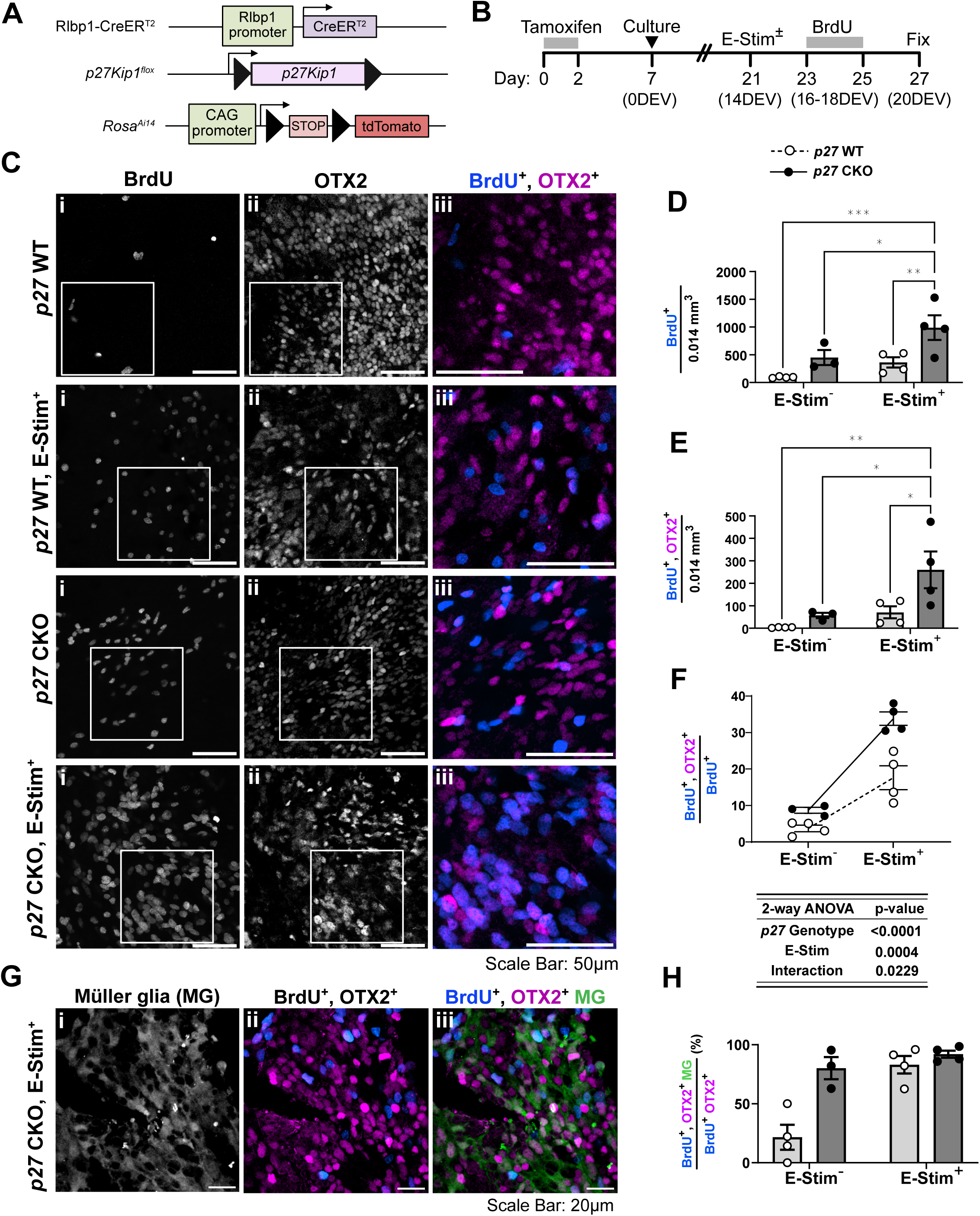
Combined *p27* inactivation and electrical stimulation synergize to induce OTX2 expression in proliferating Mueller glia. (A) Lineage reporter model with *p27^Kip^*^1^ (*p27*) inactivation driven by Mueller glia-specific Cre recombination. **(B)** Experimental timeline including tamoxifen induction of Cre recombination to inactivate *p27*, culture establishment, E-stim timepoint, BrdU treatment and fixation. **(C)** Representative 40x (80x inset) images showing BrdU and OTX2 co-labeling across four conditions (*p27* WT, *p27* WT + E-stim, *p27* CKO, *p27* CKO + E-stim). **(D,E)** Quantification of BrdU^+^ cells **(D)** and BrdU+, OTX2+ cells **(E)** per ROI volumes. **(F)** Quantification of the proportion of BrdU^+^ cells that co-express OTX2 across conditions, plotted to show individual replicates with paired E-stim^−^ vs. E-stim^+^ connections to visualize *p27* genotype vs. E-stim interaction. Statistics determined using 2-way ANOVA; p-values for *p27* genotype, E-stim, and interaction effect are shown. N=3-4. **(G)** Representative 63x image showing BrdU and OTX2 co-localization within Mueller glia (tdTomato^+^) lineage reporter in the *p27* CKO + E-stim condition. **(H)** Quantification of the proportion of BrdU^+^ OTX2^+^ cells that are tdTomato^+^, indicating Mueller glial lineage, across conditions. N=3-4.

As expected, more BrdU^+^ cells were observed in the E-stim (p27 WT, E-stim^+^) and p27 inactivation (p27 CKO) conditions compared to control (p27 WT) but, interestingly, combining p27 inactivation and E-stim caused the largest increase (p27 CKO, E-stim^+^; Figure 3C(i), D). Similar effects were observed for cells co-expressing BrdU and OTX2 (Figure 3C, E). The single conditions compared to the control were not statistically significant in this experiment because of the post-hoc correction for multiple comparisons, but the upward trends are likely true effects (Figure 3D, E). Importantly, analysis of interaction between conditions suggests a synergistic effect of the combinatorial treatment to generate BrdU^+^, OTX2^+^ cells (Figure 3F). Because of the high density of Tom^+^ cells in this experiment, it was not possible to accurately quantify BrdU and OTX2 staining in the MG-Tom population (Figure S2), but we were able to quantify the number tdTomato^+^ cells in the BrdU^+^, OTX2^+^ cell population and found that most were triple-labeled in the single and combinatorial treatments (Figure 3H,I). The untreated condition had a smaller proportion of BrdU^+^, OTX2^+^ cells that were tdTomato^+^, but this should be interpreted with caution since BrdU^+^, OTX2^+^ cells were rare in this condition (Figure 3E).

Together, these data support a model in which intrinsic p27 inactivation and extrinsic bioelectric modulation cooperate to drive proliferation-linked reprogramming in adult Mueller glia.

### Tracking proliferative Mueller glia with the H3.1 iCOUNT allele

A technical challenge to studying proliferative reprogramming is the lack of tools that can track cell type- or lineage-specific proliferation or proliferation history. Nucleotide analog labeling (i.e. BrdU, EdU) is not cell-type specific, only cells in S-phase at the time of exposure are labeled, and post-fixation staining is required for detection. Genomic barcoding can record proliferation history with lineage resolution, but these approaches are resource intensive, requiring single cell sequencing and complex barcode delivery strategies^33^. The Cre-dependent *Rosa^FUCCI^*^2a^*^R^*reporter system can identify cells in actively in cycle (S and G2 phases) with cell type- or lineage-specificity but does not reveal proliferation history^34,35^. One potential solution is the recently published H3.1-iCOUNT mouse allele (iCOUNT) that incorporates a Cre-inducible mCherry-to-eGFP switch, both of which are fused to the endogenous histone H3.1 protein (Figure 4A). H3.1 is a core histone that is synthesized and added to chromatin during S-phase and persists for the life of the cell^36^. Because of its stability, the iCOUNT allele was developed as a quantitative reporter of cell division history based on mCherry dilution and eGFP accumulation without impacting proliferation or survival (Figure 4B)^37^. We therefore assessed its utility for studying proliferative reprogramming.

**Figure 4.**
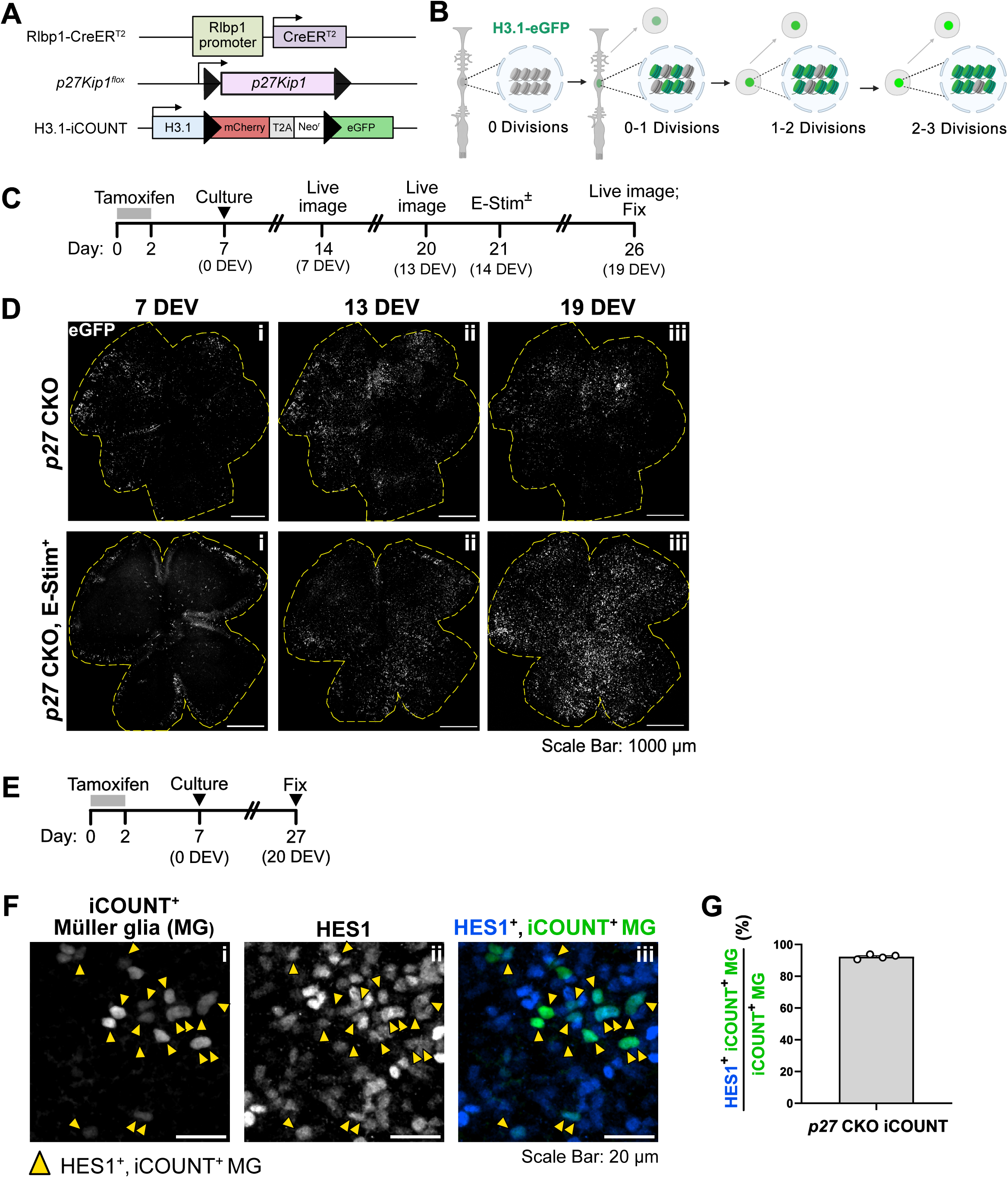
Live-cell proliferation tracking with H3.1-iCOUNT reveals increased proliferation after electrical stimulation in *p27* CKO Mueller glia. (A) Proliferation reporter model H3.1-iCOUNT and *p27* inactivation driven by Mueller glia-specific Cre recombination. **(B)** Schematic description of eGFP accumulation over successive cell divisions with the H3.1-iCOUNT model. **(C)** Longitudinal live imaging timeline and **(D)** representative 40x tiled images *of ex vivo* retinal cultures at 7 DEV, 13 DEV (pre-E-stim), and 19 DEV (post-E-stim), comparing paired *p27* CKO cultures with or without E-stim at 14 DEV, imaged on a ZEISS Axio Zoom.V16 with structured illumination to visualize iCOUNT signal. **(E)** Experimental timeline and **(F)** representative images of unstimulated *p27* CKO tissues from panel (B) showing iCOUNT signal and HES1 (Mueller glia marker) immunostaining overlap following fixation to validate Mueller glial identity of iCOUNT^+^ cells. **(G)** Quantification of the percentage of iCOUNT^+^ cells that are HES1^+^ in *p27* CKO iCOUNT retinas. N=4.

We first generated mice with iCOUNT and *Hes1^CreERT^*^*2*^ to label proliferative cells in the developing retina (Figure S3). Tamoxifen treatment at E16.5 followed by analysis at E18.5 revealed robust eGFP expression (referred to as iCOUNT^+^), confirming that the iCOUNT cassette is activated and stably incorporated into dividing retinal cells (Figure S3A-C). mCherry was difficult to detect, however, in embryonic retina by microscopy or flow cytometry (Figure S3C–D), so all subsequent analyses focused on eGFP detection rather than quantitative fluorescence intensity measurements.

Next, mice containing iCOUNT, Rlbp1-CreER^T2^, and *p27^flox^* were generated (Figure 4A). Longitudinal imaging of live tissue across the culture period (Figure 4C) revealed that *p27* CKO retinas exhibited a progressive increase in iCOUNT^+^ nuclei over time, consistent with sustained Mueller glial proliferation. E-stim treated retinas at the final timepoint show a visible increase in density of iCOUNT^+^ nuclei compared to unstimulated *p27* CKO controls (Figure 4D), again supporting enhanced accumulation of proliferative Mueller glia following E-stim. Immunostaining for HES1, which is restricted to Mueller glia in the adult retina, showed that the majority of iCOUNT^+^ cells were HES1^+^ (Figure 4E-G). These findings establish that the iCOUNT reporter can serve as a longitudinal Mueller glia-specific proliferation tracker. Together with the BrdU-based analyses in Figure 3, these results indicate that after p27 inactivation, Mueller glia sustain proliferation over days in culture that can be further enhanced by E-stim.

### p27 inactivation and E-stim bias proliferative Mueller glia toward neurogenic states

The iCOUNT reporter model provided an opportunity to specifically enrich for Mueller glia and their progeny based on proliferation history. To investigate transcriptional changes that underly the synergy between p27 inactivation and E-stim, scRNA-seq was performed on iCOUNT^+^ cells isolated from *p27* CKO *ex vivo* retinas cultured with or without E-stim (Figure 5A). p27 WT conditions were excluded from this analysis due to insufficient iCOUNT eGFP detection (Figure S4).

**Figure 5.**
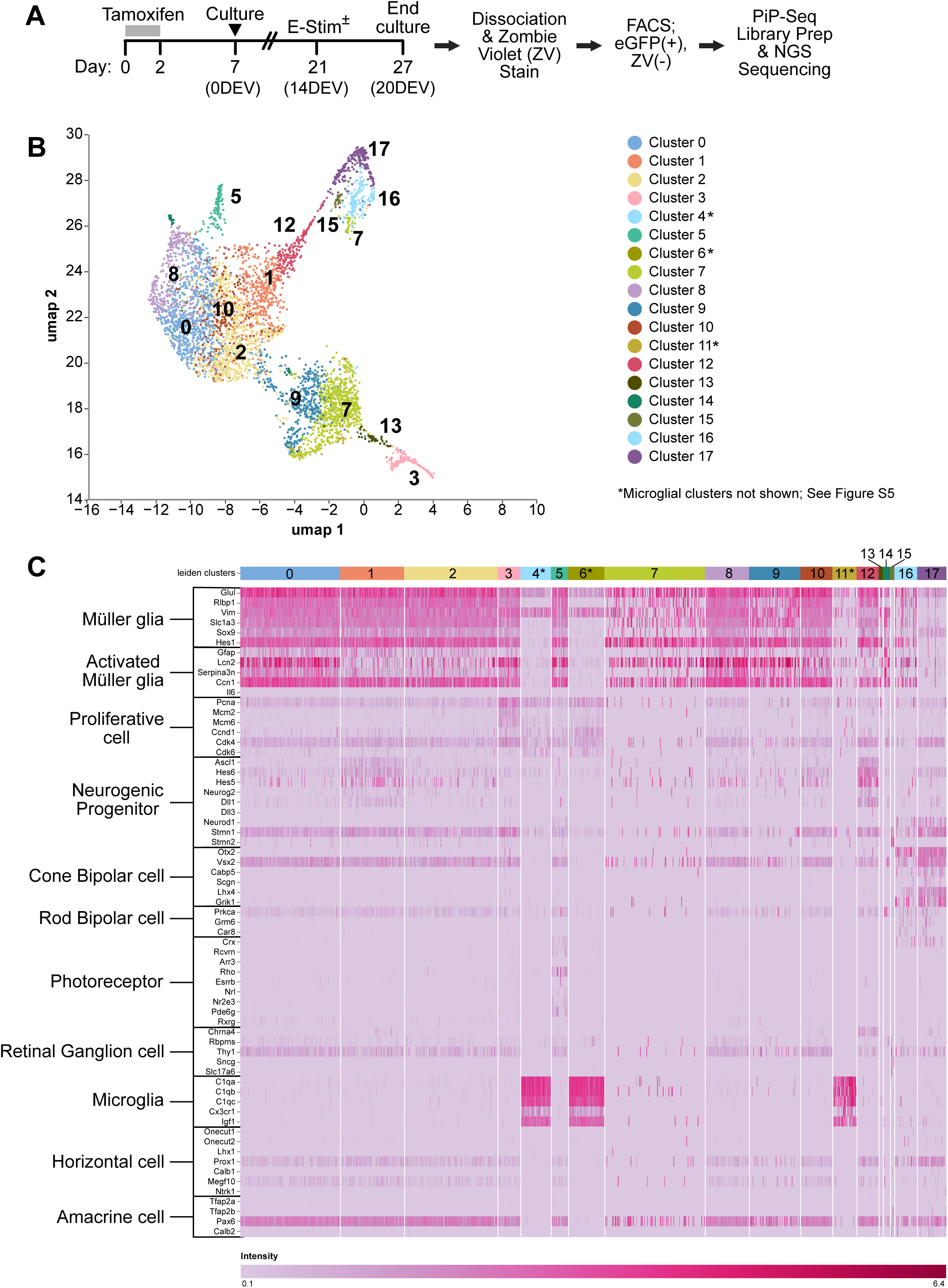
Single-cell RNA-seq workflow and cell-state annotation of iCOUNT+ cells from *p27* CKO retinas ± electrical stimulation reveals neurogenic signatures among proliferative Mueller glial populations. (A) Experimental and processing pipeline for scRNA-seq of proliferative *p27* CKO Mueller glia with or without E-stim: *ex vivo* culture of Mueller glia specific *p27* CKO with H3.1 iCOUNT retinas, E-stim or unstimulated controls at 14 DEV, dissociation at 20 DEV, Zombie Violet viability staining, FACS purification of eGFP^+^ (iCOUNT^+^), Zombie Violet− cells, and library preparation/sequencing. N=3 per condition. **(B)** UMAP visualization of Leiden clusters from integrated scRNA-seq data (microglial clusters visualized in Figure S5). **(C)** Heatmap of canonical marker gene expression used to annotate major retinal cell types and Mueller glial cell states.

Three samples per condition passed quality control, yielding 5538 cells analyzed in aggregate using a protocol optimized for small inputs (SI appendix, Table S4). Unsupervised clustering identified 17 clusters segregated into two superclusters (Figure S5). The 14-cluster group (Figure 5B) expressed genes associated with Mueller glia states, neurogenic competence, and neuronal differentiation (Figure 5C), and the 3-cluster group (Figure S5, bottom right) with microglia or immune cell identities (Figure 5C).

Cells from both conditions (p27 inactivation alone vs. combined p27 inactivation plus E-stim) were present in all clusters and global differential expression analysis did not identify markers specific to E-stim (Figure 6A, B). However, a set of clusters was preferentially enriched for E-stim^+^ cells (Figure 6C, red bars). Many of these clusters (e.g. Clusters 0,1, 8, 10) retained Mueller glial identity (*Rlbp1, Glul, Hes1*; Figure 6D,E; Figure S6A), but a distinct group of these clusters (Clusters 1, 12) was also associated with expression of neurogenic progenitor markers such as *Hes6*, *Ascl1* and *Neurog2* (Figure 6F-H). Other clusters within the E-stim enriched group (Clusters 16, 17) showed reduced expression of glial genes and upregulated expression of cone bipolar cell markers including *Otx2*, *Cabp5* and *Crx* (Figure 6I-K). These expression patterns reflect characteristics of Mueller glia passing through a neurogenic progenitor-like state and into a cone bipolar cell-like state, as observed in the ANTSi paradigm^14,15,18^. Inactivation of NFI factors (*Nfia/b/x*) has been shown to induce Ascl1 expression in non-proliferative adult mouse Mueller glia after injury^38^. These factors were not noticeably reduced in clusters associated with Ascl1 expression (Figure S6D-F), implying that Ascl1 activation with this paradigm occurs independently of NFI downregulation.

**Figure 6.**
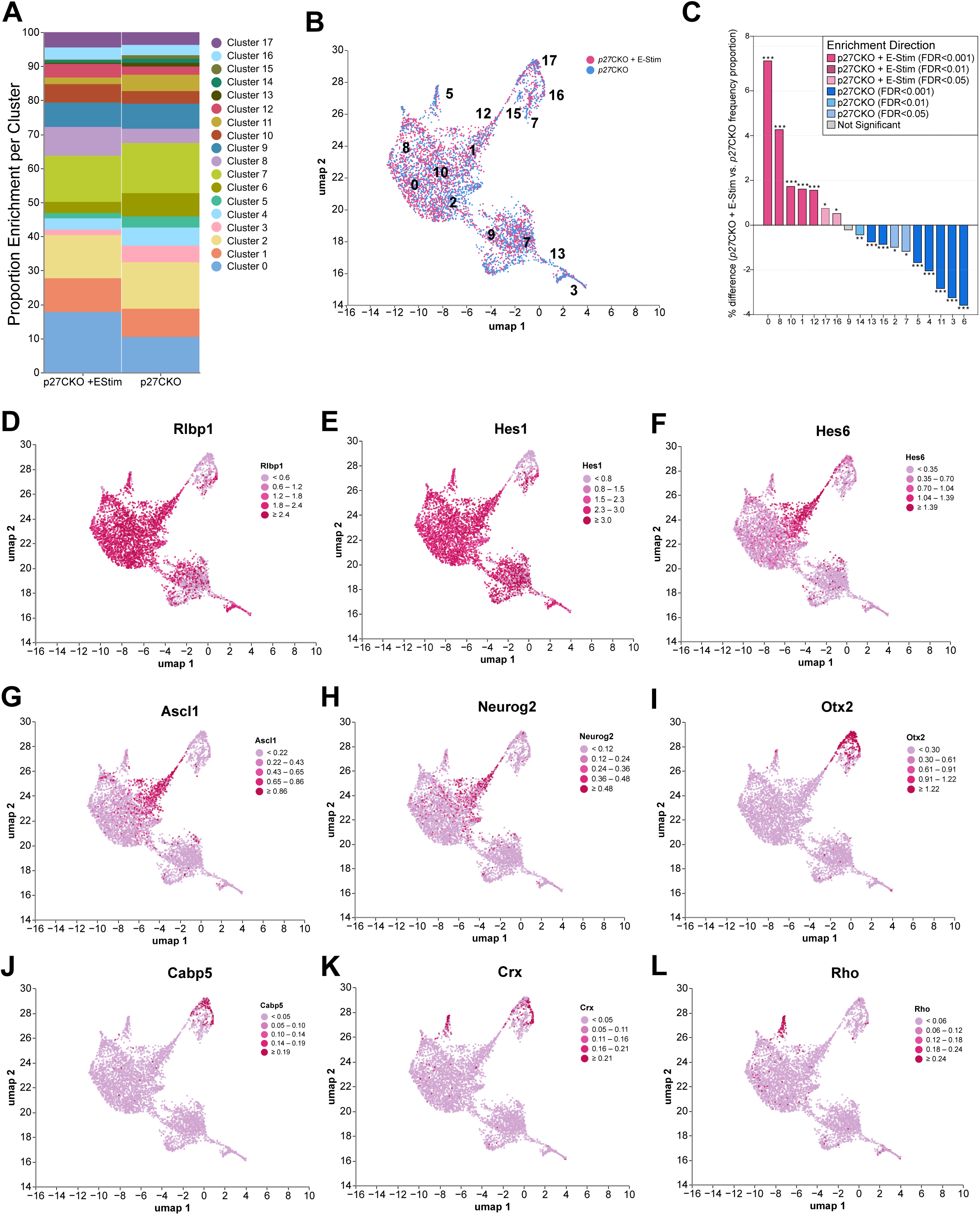
Electrically-stimulated, proliferative *p27* CKO Mueller glia are enriched into proneural and cone bipolar–like transcriptional trajectories. (A) Cluster composition by condition, shown as proportions of cells contributed by *p27* CKO vs. *p27* CKO + E-stim within each transcriptional cluster. **(B)** UMAP colored by condition (*p27* CKO vs *p27* CKO + E-stim). **(C)** Cluster enrichment analysis shows direction and significance of condition-associated enrichment across clusters. Statistics analyzed with Fisher’s exact test, FDR thresholds as indicated. **(D,E)** Feature plots for Mueller glial markers (*Rlbp1, Hes1*) across the UMAP embedding. **(F–H)** Feature plots for proneural/neurogenic markers (*Hes6, Ascl1, Neurog2*). **(I,J)** Feature plots for cone bipolar-associated markers (*Otx2, Cabp5*). **(K)** Feature plots for bipolar- and photoreceptor-associated marker *Crx*. **(L)** Feature plot for rod photoreceptor-associated gene *Rho,* highlighting a distinct, low frequency trajectory enriched in unstimulated *p27* CKO cells.

In contrast, a second set of clusters (3, 7, 9, 13) is enriched for E-stim^−^ cells (Figure 6A–C; blue bars in C). These clusters show expression profiles consistent with Mueller glial identity (Rlbp1, Hes1, Glul) with some cells in these clusters (7, 13) exhibiting reduced NFI factor expression (*Nfia/b/x*; Figure S6D-F). However, this does not resolve in increased progenitor marker expression as reported in NFI inhibition studies^6,38^. Interestingly, these clusters show reduced expression of genes known to be expressed in adult Mueller glia that also play important roles in progenitor differentiation and neurogenesis during retinal development (Pax6, Vsx2; Figure S6B, C). A subset of cluster 7 is associated with clusters containing cells demonstrating bipolar cell-like expression, but these cells show reduced *Otx2*, *Cabp5* and *Crx* expression compared to their neighbors (Figure 6I-K). These findings support prior findings *in vivo* that in the absence of E-stim, p27-deficient Mueller glia are more likely to retain glial properties, consistent with prior findings *in vivo*^26,22,30^.

Additionally, a small but distinct cluster that is enriched for p27-deficient Mueller glia without E-stim co-expresses glial genes and rod photoreceptor-associated genes including *Rho*, *Pde6g*, and *Nr2e3* (Figure 6L; Figure S6G,H). This cluster may represent a rare, alternative cell state in which p27 inactivation alone initiates partial neurogenic reprogramming of Mueller glia toward a rod photoreceptor-like state, a suggestion that has been controversial across *in vivo* studies^31,32^. It is unclear from this data if photoreceptor gene expression in this population is due to activation of photoreceptor differentiation pathways or if this is a consequence of trogocytosis or incomplete lysosomal degradation after phagocytosis^39,40^. Regardless, the presence of this cell population was low compared to the clusters expressing bipolar cell genes, supporting a neurogenic conversion toward the cone bipolar–like state after p27 inactivation and E-stim.

To validate key scRNA-seq findings at the protein level, p27 inactivation and E-stim were repeated with iCOUNT retinas and assessed for proliferation and reprogramming (Figure 7A, B). Consistent with our observations using BrdU, p27 inactivation combined with E-stim increased the number of iCOUNT^+^ nuclei over p27 inactivation alone, indicating accumulation of proliferative Mueller-derived cells (Figure 7C(i), D). Additionally, p27 inactivation combined with E-stim significantly increased ASCL1-expressing cells (Figure 7C(ii), E), with a greater fraction of iCOUNT+ cells positive for ASCL1 (Figure 7C(v), F). The window of BrdU treatment in this experiment was expanded to capture all cell cycle re-entry following E-stim. Many ASCL1^+^ Mueller glia were BrdU^+^, indicating activation of proneural signals within an actively proliferative population (Figure 7C(iv, vi), G). Similarly, a significant fraction of iCOUNT^+^ cells expressed OTX2 and CABP5 following E-stim (Figure 7H-K), indicating that E-stim increases the efficiency with which proliferative Mueller-derived cells undergo neuronal conversion.

**Figure 7.**
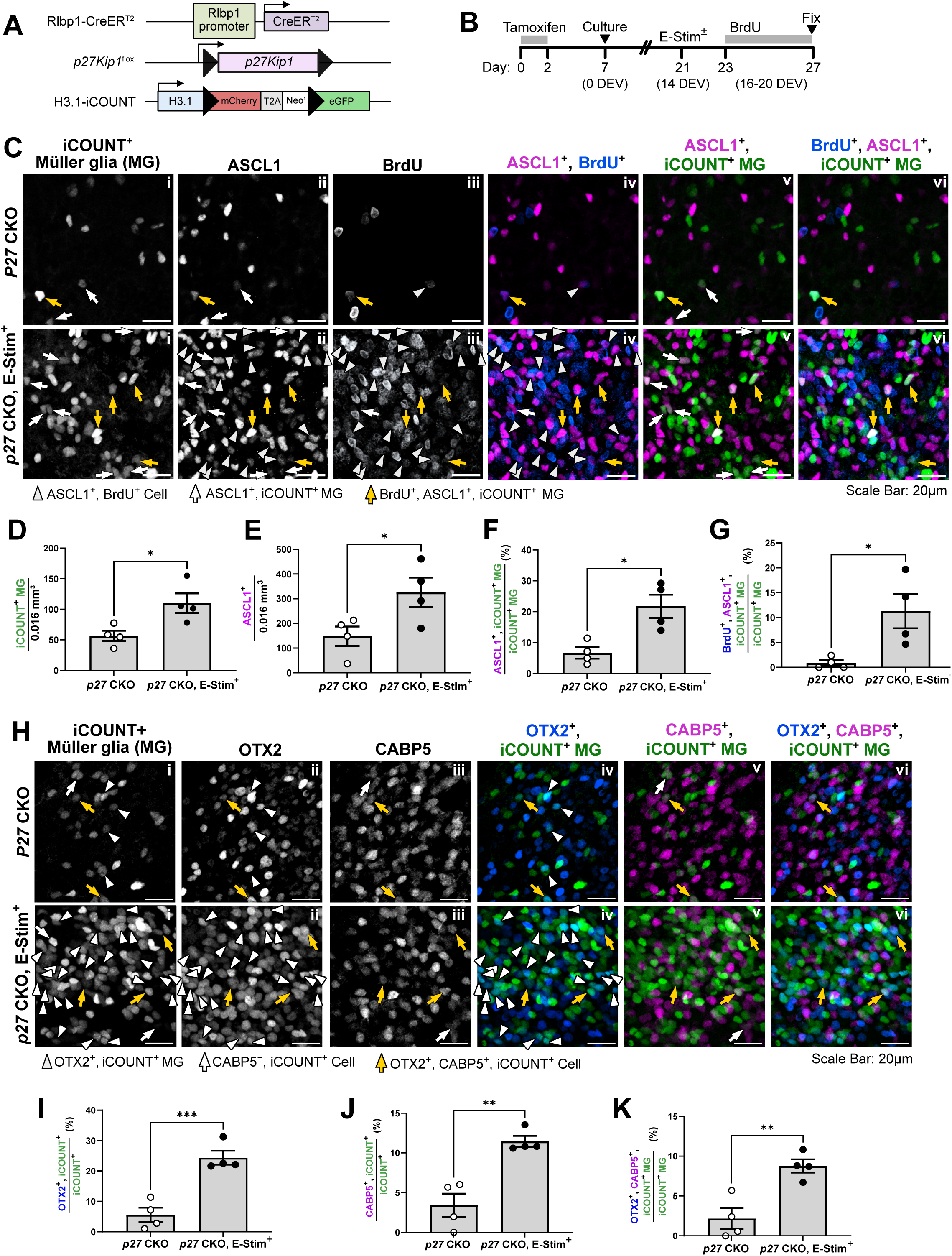
Protein-level validation of scRNA-seq reveals proneural and bipolar cell-associated marker expression in proliferative *p27* CKO Mueller glia following electrical stimulation. (A) Proliferation reporter model H3.1-iCOUNT and *p27* inactivation driven by Mueller glia-specific Cre recombination. **(B)** Experimental timeline including tamoxifen activation, culture establishment, E-stim, and extended BrdU treatment period to capture all proliferative cells following E-stim. (**C)** Representative 40x images of whole-mount retinas showing iCOUNT^+^ *p27* CKO Mueller glia with ASCL1 immunostaining to identify neurogenic competence and BrdU labeling to show active proliferation following E-stim. **(D–G)** Quantification of iCOUNT^+^ Mueller glia 0.016 mm^3^ ROI volume **(D)**, ASCL1^+^ cells per 0.016 mm^3^ ROI volume **(E)**, percentage of iCOUNT^+^ Mueller glia that are ASCL1^+^ **(F)**, and percentage of iCOUNT^+^ Mueller glia that are BrdU^+^ and ASCL1^+^ **(G)** in *p27* CKO vs. *p27* CKO + E-stim retinas. Statistics in all panels analyzed using Student’s T-test. N=4. **(H)** Representative 40x images showing iCOUNT^+^ Mueller glia co-stained for OTX2 and CABP5. **(I–K)** Quantification of the percentage of iCOUNT^+^ Mueller glia that are OTX2^+^ **(I)**, CABP5^+^ **(J)**, or OTX2^+^, CABP5^+^ **(K)** in *p27* CKO vs. *p27* CKO + E-stim retinas. N=4.

To assess if these changes were accompanied by increased apoptosis, expression of apoptosis marker cleaved-CASP3 was examined. CASP3^+^ cells were sparse across conditions, and no qualitative increase in CASP3 labeling of iCOUNT^+^ cells was detected following E-stim (Figure S7). This suggests that apoptosis is unlikely to be a major driver of proliferation or reprogramming after E-stim.

## Discussion

Promoting robust proliferation, loss of Mueller identity, and activation of neural differentiation programs are complex, sometimes competing processes that likely require precise timing and coordinated manipulations. Using 3D adult mouse retinal culture as a tissue-context testing platform, we found that combined electrical stimulation and Mueller glia-specific p27 inactivation, which we refer to as ESPI, and show how these inputs cooperate to couple cell cycle reentry with neurogenic progression in Mueller glia. E-stim alone increased the proportion of Mueller glia that re-entered the cell cycle and induced endogenous ASCL1 and OTX2 expression, but proliferation and neurogenic competence were uncoupled with minimal evidence of cells adopting neuronal fates. p27 inactivation alone generated a large proliferative Mueller-derived population, yet by itself did not reliably drive neurogenic progression. ESPI shifted proliferative Mueller-derived cells into neurogenic-competent states at higher frequency and was associated with acquisition of bipolar-associated differentiation markers, identifying ESPI as an effective strategy to achieve neural differentiation of Mueller-derived cells in adult retinal tissue.

Prior literature supports the premise that bioelectric cues can influence glial behavior and neurogenic programs. In the retina, E-stim of various parameters has been explored most prominently for neuroprotection and functional preservation^41–45^. Recent studies show that E-stim promotes long-distance axon regeneration that partially restores vision in adult rats after optic nerve crush injury^46,47^. Electrical stimulation in 2D culture of enriched Mueller glia from postnatal mouse retinal tissue (P6-7) reported changes in Mueller glial activation, proliferation, and gene expression after stimulation^48,49^. In neural stem cells and fibroblasts, electrical fields can bias neural stem or progenitor behavior and promote neuronal differentiation programs^50–53^.

The induction of endogenous Ascl1 in this study is particularly notable because adult mammalian Mueller glia typically do not upregulate Ascl1 following injury^18,19,54,55^. This contrasts strongly with regenerative species such as zebrafish, where injury induces Ascl1a in Mueller glia and is required for dedifferentiation and progenitor formation^3,56,57^. Our observation that E-stim can activate endogenous Ascl1 in adult Mueller lineage cells suggests that bioelectric modulation can engage a proneural entry point that mammalian Mueller glia normally fail to access following injury. E-stim alone, however, might not be sufficient for neurogenic-competent cells to progress to specific neuronal fates.

Prior work demonstrates that p27 is required to maintain Mueller glia in non-proliferative, non-reactive state, but evidence suggests a role for p27 inactivation in cell reprogramming as well. p27 has been shown to directly repress Sox2 expression during exit from pluripotency, and that inducing pluripotency with the Yamanaka factors in p27-inactivated fibroblasts does not require Sox2 overexpression^58,59^. Additionally, Notch activation influences Mueller glial plasticity, evidenced by latent stem cell-like gene expression that coincides with reduced p27 expression through Skp2-mediated degradation^29,60,61^. Our study advances this framework by providing evidence that p27 inactivation can support an intermediate progenitor-like fate when paired with an instructive cue. Within an enriched population of proliferative, p27-inactivated Mueller-derived cells, scRNA-seq identifies clusters expressing proneural/progenitor-associated markers (*Hes6*, *Ascl1*, *Neurog2*) that transition toward Otx2/Cabp5-expressing bipolar-associated states, and these trajectories are preferentially enriched by E-stim. This intermediate state is important because it represents a transitional window in which fate restriction appears partially relieved (glial programs reduced, neurogenic competence increased), creating an opportunity for testing signals that promote different cell fates, a major goal in Mueller glial reprogramming to restore tissue function.

One possible mechanism by which ESPI promotes these transitions is that E-stim intersects with Notch-associated repression of proneural programs. Adult Mueller glia express the Notch effector Hes1^62,63^, and sustained Notch signaling has been implicated as a barrier to durable neuronal conversion^29,6^. Enrichment of Hes6 in the E-stim-enriched population, alongside activation of Ascl1 expression, is consistent with relief of Hes1-like repression, allowing stabilization of proneural programs and progression toward differentiation^18,64,65^.

Despite persistent interest in Mueller glial reprogramming, neurogenic outcomes remain relatively rare, and a promising path forward is combinatorial intervention^6,14,15,66–68^. 3D *ex vivo* retina culture provides a practical framework to address feasibility issues inherent to iterative testing *in vivo*, but comes with its own limitations, such as neuronal degeneration, alterations in vascular, systemic, and immune cues, and prolonged culture imposes metabolic and stress constraints that could shift glial competence^8,11,69^. These differences could contribute to the rare appearance of ASCL1^+^ cells observed in unstimulated cultures. Additionally, the current state of 3D culture methods are unlikely to be sufficient for studies analyzing long term tissue restoration or incorporation of new neurons into existing circuitry. Future studies aimed at culture system improvement to promote long term survival of all retinal cell types will help to advance research into reprogramming and regeneration^8^. Still, this model provides a robust platform to test first tier outcomes of candidate reprogramming strategies, and differences in *ex vivo* and *in vivo* outcomes provide an opportunity to identify environmental barriers in the intact retina that limit Mueller glial reprogramming and allows prioritization of interventions that reduce those constraints.

This study incorporates the H3.1-iCOUNT allele to enable Mueller glial lineage tracing and proliferation-history tracking in one reporter. Coupled with nucleotide analog labeling, iCOUNT allows stimulus-specific proliferative responses to be mapped longitudinally and enables enrichment of cells based on proliferation status, approaches that have been lacking in reprogramming studies. A reported limitation of this model is that iCOUNT can exhibit mosaicism of activation beyond what is expected with cre recombination, raising the possibility that proliferation may be underreported^37^. Accordingly, iCOUNT-based analyses should be interpreted within the context of the iCOUNT+ population. This is similar to nucleotide analog labeling where a label-negative cells does not rule out proliferation outside the labeling interval. Even with the limitation of mosaicism, the iCOUNT reporter system provides a new context for studying reprogramming within proliferative cell populations that could improve efficiency and interpretation of reprogramming outcomes.

In summary, ESPI pairs extrinsic bioelectric modulation with intrinsic cell-cycle release to increase the frequency proliferative reprogramming in Mueller-derived cells. This study suggests that p27 loss creates a proliferative context characterized by increased plasticity and instability of differentiated glial identity, and E-stim biases this plasticity toward endogenous Ascl1 activation and progression toward bipolar-associated differentiation programs. More broadly, ESPI and the *ex vivo* framework provide an accessible route to systematically discover and optimize combinatorial strategies that couple proliferation to neurogenic competence in adult mammalian retina.

## Materials and Methods

Extended methods pertaining to animal care, mouse strains, tamoxifen administration, dissection, culture establishment and maintenance, drug delivery, fixation and ScRNA-seq preparation and analysis are provided in SI appendix. Supplemental tables list information on: (S1) Key resources, (S2) Electrical stimulation parameters and setup, (S3) Antibodies and working dilutions, and (S4) scRNA-seq sample metadata and QC summary.

### Experimental design

Adult mouse retinas were maintained as *ex vivo* 3D whole-retina cultures at the air-liquid interface as previously described^13^ (see SI Appendix, Extended Methods). Days *ex vivo* (DEV) denotes days in culture (0 DEV = day of culture start). Where indicated, paired retinas from the same animal were assigned to stimulated versus unstimulated conditions to control for inter-animal variability. Genetic models used include Mueller glia lineage tracing, conditional p27Kip1 inactivation, and the H3.1-iCOUNT proliferation-history reporter (see SI Appendix, Extended Methods and Table S1).

### Agarose-hydrogel mediated electrical stimulation (E-stim)

E-stim was delivered to 3D cultures at 14 DEV using the agarose dish electroporation setup as described previously^13^. Stimulation consisted of one treatment of five square-wave pulses (0.0417 kV/cm (Voltage = 25 V, distance = ∼0.6cm), 50 ms each) with 250 ms intervals (alternating current) with PBS & loading dye present in electrical stimulation solution. Electrode geometry, spacing/field strength, and all device settings are provided in SI Appendix, Table S2.

### Whole retinal piece immunohistochemistry

Retinal tissue was separated into 4 pieces to maximize staining per tissue, but each N per experiment came from separate tissues. Incubation times for blocking, washes, and antibody exposure were extended to maximize penetration (see SI Appendix, Extended Methods). Product information on antibodies used is available in Table S3.

For BrdU co-staining, other markers were stained first and tissues were post-fixed for 30 minutes in 4% PFA prior to BrdU antigen retrieval with 2 N HCl (45 minutes, room temperature) followed by neutralization in 0.1 M sodium borate (20 minutes).

### Imaging

Longitudinal live imaging of iCOUNT fluorescence was performed using wide-field epifluorescence (ZEISS Axio Zoom.V16 with Apotome 3), and fixed-tissue imaging was performed on a ZEISS LSM 710 confocal microscope. Image processing was performed in FIJI/ImageJ (see SI Appendix, Table S1).

### Single cell preparation and scRNA-seq

At 20 DEV, retinal cultures were dissociated using papain, stained with Zombie Violet viability dye, and viable iCOUNT-positive (eGFP-positive) cells were enriched by fluorescence-activated cell sorting (Cytek Aurora CS). Single-cell libraries were prepared using the PIPseq V-T2 3’ single-cell kit. Reads were quantified with the PIPseeker V pipeline using a custom reference that included eGFP, and downstream analysis was performed in Trailmaker with batch integration by Harmony and doublet detection by DoubletFinder (SI Appendix, Extended Methods, Tables S1, S4).

### Quantification and statistics

Image-based quantification was performed on blinded confocal z-stacks using consistent thresholds within experiments and Imaris for segmentation and colocalization. Each n represents one retina (no more than one retina per mouse per condition). Two-group comparisons were analyzed using paired two-tailed Student’s t-tests, and multi-group comparisons were analyzed using two-way ANOVA with Tukey post hoc tests. Significance was set at p ≤ 0.05.

## Acknowledgements

Funding for this study was provided by the National Eye Institute (R01-EY013760; R21-EY033473; P30-EY008126), the Janet and Jim Ayers Research Fund in Regenerative Visual Science, the William A. Black Chair in Ophthalmology, a Potocsnak Discovery Grant in Regenerative Neuroscience, and an unrestricted grant from Research to Prevent Blindness, Inc. M.L.S was supported by the Vanderbilt Vision Training Grant (T32-EY007135) and a Ruth L. Kirschstein Predoctoral Individual National Research Service Award (F31-EY035554) from the National Eye Institute. FACS sorting and analysis were performed by Christian Warren at the VUMC Department of Veterans Affairs Flow Cytometry Core. Confocal imaging and Imaris software analysis were performed at the Vanderbilt Cell Imaging and Shared Resource. The Zeiss LSM710 confocal microscope used for this study was acquired with funding from NIH (S10-RR027396). Access to a Zeiss AxioZoom V16.Apotome 3 imaging system was generously provided by Dr. Ian Macara (Vanderbilt University). We thank Dr. Thomas Reh (University of Washington) for providing the ASCL1-overexpression mouse model. We also thank Francesca Napoli for assistance in retinal culture preparation, and members of the Levine and Fuhrmann laboratories, and M.L.S thesis committee for their insights and feedback.

## Author Contributions

Conceptualization, E.M.L. and M.L.S.; methodology E.M.L., J.J., and M.L.S.; formal analysis, M.L.S.; investigation, M.L.S.; resources, E.M.L.; data curation, M.L.S.; writing (original draft), M.L.S.; writing (review and editing), M.L.S., J.J., and E.M.L.; visualization, M.L.S.; supervision, E.M.L.; project administration, E.M.L.; funding acquisition, E.M.L.

