## Supplemental Information for "Electrical stimulation combined with p27Kip1 inactivation drives proliferative neurogenic reprogramming of Mueller glia in the adult mouse retina"

This file contains: Extended methods, Supplemental figures and figure legends

**EXTENDED METHODS**

**Animal care**

Adult mice (7–12 weeks of age; female and male) were used, housed on a 12h/12h light-dark cycle with food and water *ad libitum*. All transgenic mice are backcrossed to B6J/129 mice. All procedures were conducted under Vanderbilt Institutional Animal Care and Use Committee approval, following ARVO guidelines and in concordance with NIH guidelines for Responsible Conduct in Research (RCR). All possible efforts were made to minimize animal suffering and the number of animals used.

**Mouse Strains**

GlastCreER; LNL-tTA; teto-mAscl1-ires-eGFP; CC-eGFP (Tg(Slc1a3-cre/ERT)1Nat/J (JAX: 012586); B6.129P2(Cg)-$Gt(ROSA)26Sor^{tm1(tTA)Roos}$/J (JAX:011008); M. Nakafuku, U. Cincinnati; FVB.B6-Tg(CAG-cat,-EGFP)1Rbns/KrnzJ (JAX: 024636)). Provided as a gift from the laboratory of Dr. Thomas Reh (University of Washington, Seattle). 7-12 weeks old, female and male. Used for experiments in figure 1.

Rlbp1-$CreER^{T2}$; $Rosa^{Ai14}$ (Tg(Rlbp1-cre/ERT2)1Eml (MGI:7708085)); B6.Cg-Gt(ROSA)26Sortm14(CAG-tdTomato)Hze/J (JAX: 007914)). Used for experiments in figures 1, 2, 3, and S1.

$Cdkn1b^{flox/flox}$(129-$Cdkn1b^{tm1Mlf}$/J (JAX: 003122)). Crossed with Rlbp1-$CreER^{T2}$; $Rosa^{Ai14}$ for experiments in figures 3 and S2.

H3.1-iCOUNT ($H3c6^{em1Sjes}$ (JAX: 037318)). Crossed with Rlbp1-$CreER^{T2}$ in figure S4 and Rlbp1-$CreER^{T2}$; $Cdkn1b^{flox/flox}$ in figures 4-7, S3, and S5-S7.

Hes1:$CreER^{T2}$ ($Hes1^{tm1(cre/ERT2)Lcm}$ (MGI: 4412375)). Crossed with H3.1 iCOUNT in figure S3.

**Tamoxifen administration**

All experiments used adult mice aged 7-12 weeks. Cre recombination for experiments described in figures 2-7 and all supplemental figures was induced by oral gavage of tamoxifen dissolved in corn oil. For experiments described in figure 1, Cre recombination was induced via gel food mixed with tamoxifen dissolved in corn oil as recently described[37]. Tamoxifen administration to mice from figure 1 followed dosing reported in the original ANTSi studies[5,6]. In all other experiments, Cre recombination in adult Müller glia was induced by oral gavage of tamoxifen in corn oil using the high-dose paradigm previously described (2 doses, 200 μg/gram body weight). Retinas were harvested 5-7 days after initial tamoxifen induction.

**Retinal Dissection**

Retinas were dissected as described in previous work from this lab[11] in cold HBSS (with $Ca^{2+}$/$Mg^{2+}$) supplemented with HEPES and antibiotics/antimycotics. Tools and work area sterilized before tissue collection. Eyes were enucleated, anterior segments removed, and the neural retina was isolated with the optic nerve head retained for orientation.

**3D whole retinal culture**

3D whole retinal cultures were established at the air–liquid interface using the culture system described in our previous study[11]. "Days *ex vivo*" (DEV) refers to the number of days after placement into culture (0 DEV = day of culture start), consistent with the experimental timelines shown in the figures (e.g., E-Stim at 14 DEV).

**Drug delivery to 3D culture**

All pharmacologic treatments and BrdU labeling were delivered directly into culture medium with a drop of media containing drug put on top of the tissue daily. Vehicle controls received an equivalent final concentration of solvent. Prior studies validated the ability for BrdU to be taken up into the tissue when delivered in culture medium[11]. In experiments for figure 1, 16µM BrdU and (where indicated) small-molecule treatments were applied beginning at 2 days *ex vivo* (DEV) and maintained through 4 DEV (48 h total), followed by a wash out and continued culture periods without drug until fixation at 16 DEV, in accordance with methods described in the original ANTSi studies[5,6]

Trichostatin A (TSA) and the STAT inhibitor SH-4-54 were delivered only for experimental conditions in figure 1, via the culture medium beginning at 2 DEV for 48 h (2–4 DEV), after which culture medium was replaced and cultures continued without drug until fixation. Stock solutions were prepared in DMSO consistent with prior ANTSi implementations (e.g., TSA prepared in DMSO at 1 µg/µL; SH-4-54 prepared at 10 mM in TSA-containing DMSO) and diluted into culture medium to the working concentrations used for each experiment.

In all other applicable experiments, 16µM BrdU was added to culture medium as indicated during defined labeling windows that varied by experiment. In E-Stim experiments, BrdU was added after stimulation during a defined post-stimulation window (1-2 DEV) to assess prolonged proliferation after the initial injury response and to avoid BrdU incorporation that may occur due to extracellular influx immediately following E-Stim.

**Whole tissue fixation**

Culture medium was removed and replaced with room temperature PBS to wash 3 times for 10 minutes each. 100µL 1X PBS was gently pipetted on top of tissue for each wash so as not to dislodge the tissue from the filter. Retinas were fixed in 4% paraformaldehyde (PFA) for 2 hours at ambient temperature, then overnight at 4°C. PFA was removed and fixed retinas were washed 3 times with 1 mL 1X PBS for 30 minutes at ambient temperature. Fixed retinas were stored still attached to the filters at 4°C in 1 mL 1X PBS with 0.01% sodium azide (Sigma Aldrich, 08591).

**Live 3D culture imaging**

Longitudinal imaging of whole retina volumes to monitor iCOUNT expression over time in culture (Figure 4B) was done with wide-field epifluorescence using a Zeiss Axio Zoom.V16 equipped with Apotome 3 structured illumination. Image volumes were post-processed using Apotome 3, followed by stitching with Zen 2.3 software. Images shown in Figure 4B are maximum intensity projections generated with FIJI version 2.9.0.

**Fixed tissue imaging**

All other images were captured after fixation alone or fixation and immunostaining using the Zeiss LSM 710 confocal microscope after fixation alone or fixation and staining. Images were analyzed using FIJI version 2.9.0.

**Quantification and statistical analysis of immunostaining data**

***(incorporate into text here or in ScRNA seq section): Given the variability inherent to low-input single-cell workflows, condition-associated shifts were identified by enrichment patterns, or the relative proportions of cells from each condition within each transcriptional cluster.*

All cell counting and tissue volume calculations were calculated from blinded 40X, Z-Stack, confocal image sets using consistent thresholds across matched conditions within an experiment. Segmentation, quanitifcation, and colocalization were performed using Imaris 10.2.0 software. Statistical tests (performed in GraphPad Prism 10), findings and number of samples (n) are described in each figure and associated legend. Each N refers to one retina. No more than one retina per mouse was used per condition. Comparisons between two groups were made using paired two-tailed Student's T-tests. Comparisons between more than two groups were made by two-way ANOVA with Tukey post-hoc test. Differences were considered statistically significant at $p \leq 0.05$. Data are presented as mean ± SEM. Statistical significance is indicated with asterisks: $*p \leq 0.05$, $**p \leq 0.01$, $***p \leq 0.001$, $****p \leq 0.0001$. Figure graphics created using Biorender.

**Retinal dissociation, viability staining, and FACS enrichment for ScRNA-Seq**

Retinal cultures were dissociated using the Worthington Papain Dissociation System (LK003150). At the end of the culture period, retinas were transferred to ice-cold HBSS containing $Ca^{2+}/Mg^{2+}$ supplemented with HEPES and glucose ("HBSS+"; 20 mM HEPES, ~0.6% glucose), washed, and incubated in freshly activated papain prepared in EBSS (with phenol red) and supplemented with DNase I. Samples were digested at 37 °C with gentle rotation (~30 rpm) for ~15–20 min until tissue fragmented, then triturated (15–20 passes) to generate a single-cell suspension. Enzymatic activity was quenched by addition of ovomucoid inhibitor solution supplemented with DNase I, followed by gentle mixing. The suspension was passed through a 40 µm filter and resuspended in $Ca^{2+}/Mg^{2+}$-free HBSS containing 1% BSA and RNase inhibitor (0.2 U/µL) for sorting. Cells were stained with Zombie Violet Fixable Viability Dye (BioLegend 423113; 1:1000) for 20 min at room temperature in the dark, washed once (300 × g, 10 min), and resuspended in Ca/Mg-free HBSS containing 1% BSA and RNase inhibitor prior to sorting.

Viable (Zombie Violet$^-$), iCOUNT$^+$ (eGFP$^+$ ) cells were isolated via fluorescence-activated cell sorting (FACS) using a Cytek Aurora CS (serial S0127) at the Vanderbilt Veteran's Affairs Flow Cytometry Core. Events were gated to exclude debris (FSC/SSC), select singlets, and exclude dead cells (Zombie Violet$^-$), and the target population was enriched based on GFP fluorescence (with mCherry recorded as an additional fluorescence parameter). Sorting was performed in Purity mode using a 100-µm nozzle at 18.5 psi.

**PiP-seq library preparation and sequencing**

FACS purified iCOUNT$^+$ cells for 8 samples (N=4/condition) were pelleted (300 × g, 10 min) and resuspended in the manufacturer-provided cell suspension buffer prior to library preparation. Single-cell RNA-seq libraries were generated using the PIPseq V-T2 3′ Single Cell RNA Kit (Fluent BioSciences, PIPseq-V-T2-3) according to the manufacturer's protocol, including partition-based capture, reverse transcription, cDNA amplification, and dual-index sample index PCR. Final libraries were quantified and quality-checked prior to pooling and sequencing on an Illumina platform. Libraries were sequenced using the kit-recommended read structure: Read 1 ≥45 bp (cell barcode/UMI), dual 10-bp i7 and i5 indices, and Read 2 ≥72 bp (cDNA insert). Target sequencing depth followed manufacturer guidance (starting at ~20,000 reads per cell). One control condition sample was excluded during QC due to low fragment size & RIN.

**scRNA-seq quality control, integration, and clustering**

Base call files were converted to FASTQ files and reads were quantified to gene-by-cell count matrices using the PiPseeker V pipeline. To enable detection of the reporter transcript, a custom reference was generated by appending the eGFP sequence to the GRCm39 mouse reference prior to read quantification. Count matrices were processed in Trailmaker for quality control, integration, and downstream analysis. The processed Seurat object is available as a downloadable resource (**TBD**) and raw reads are available at **TBD**. Cells were filtered using standard scRNA-seq QC metrics (including transcript/UMI counts, detected genes, and mitochondrial transcript fraction), and putative doublets and ambient RNA contributions were addressed using DoubletFinder prior to downstream analyses. One experimental condition sample was excluded during QC metric filtering due to high mitochondrial transcript, resulting in a final N=3 per condition. Biological replicates were integrated with Harmony to mitigate batch effects, followed by dimensionality reduction (PCA), graph-based clustering, and visualization with UMAP. Cluster identities were assigned using canonical marker genes assembled from marker gene lists published by the Blackshaw lab at Johns Hopkins University[35], and differential expression testing. Visualizations were generated using Trailmaker and enrichment analysis statistics were calculated with Fisher's exact test.

**Quantification & statistics**

All cell counting and tissue volumes were calculated from blinded 40X, Z-Stack, confocal image sets using consistent thresholds across matched conditions within an experiment. Segmentation, quantification, and colocalization were performed using Imaris. Statistical tests (performed in GraphPad Prism 10), findings and number of samples (n) are described in each figure and associated legend. Each N refers to one retina. No more than one retina per mouse was used per condition. Comparisons between two groups were made using paired two-tailed Student's T-tests. Comparisons between more than two groups were made by two-way ANOVA with Tukey post-hoc test. Differences were considered statistically significant at $p \leq 0.05$. Data are presented as mean ± SEM. Statistical significance is indicated with asterisks: $*p \leq 0.05$, $**p \leq 0.01$, $***p \leq 0.001$, $****p \leq 0.0001$. Figure graphics created using Biorender.

**Table S1. Key resources.**

| MATERIALS AND RESOURCES | SOURCE | IDENTIFIER |
|---|---|---|
| **Chemicals and Reagents** | | |
| Agarose | RPI | A20090 |
| Antibiotic-Antimycotic | Gibco | 15240062 |
| BrainPhys Neuronal Medium | STEMCELL Technologies | 05790 |
| NeuroCult Plating Medium | STEMCELL Technologies | 05713 |
| SM1 Neuronal Supplement | STEMCELL Technologies | 05711 |
| N2 Supplement-A | STEMCELL Technologies | 07152 |
| Penicillin-Streptomycin | ThermoFisher | 15140 |
| L-Glutamine | Gibco | 25030149 |
| HBSS with Ca2+ and Mg2+ | Gibco | 14025 |
| PBS | Corning | 21-040-CV |
| HEPES | Sigma-Aldrich | H0887 |
| Triton X-100 | Thermo Scientific | A16046 |
| Normal donkey serum | SouthernBiotech | 003001 |
| Paraformaldehyde | Fisher Scientific | 50-980-495 |
| Sodium azide | Sigma-Aldrich | 08591 |

| | | |
|---|---|---|
| Hydrochloric acid (2 N for BrdU retrieval) | Fisher Scientific | SA541 |
| Sodium borate | MilliporeSigma | 1066690010 |
| Fluoromount-G | Invitrogen | 00-4958-02 |
| Glycerol | Sigma-Aldrich | G7893 |
| Methyl green | MCE | HYD0163 |
| Ethanol | Fisher Scientific | 04355223 |
| Corn oil | Sigma-Aldrich | C8267 |
| Tamoxifen | Sigma-Aldrich | T2859 |
| BrdU | Invitrogen | B23151 |
| Trichostatin A (TSA) | Selleck Chem | S1045 |
| SH-4-54 | Selleck Chem | S7337 |
| DMSO | Invitrogen | D12345 |
| **Commercial Assays** | | |
| Worthington Papain Dissociation System | Worthington | LK003150 |
| Zombie Violet Fixable Viability Dye | BioLegend | 423113 |
| PIPseq V-T2 3' Single Cell RNA Kit | Fluent BioSciences | V5.0 |
| **Experimental models: Organisms/strains** | | |
| Mouse: Tg(Slc1a3-cre/ERT)1Nat/J (GlastCreER) | The Jackson Laboratory | JAX:012586 |
| Mouse: B6.129P2(Cg)-Gt(ROSA)26Sortm1(tTA)Roos/J (Rosa-LNL-tTA) | The Jackson Laboratory | JAX:011008 |
| Mouse: teto-mAscl1-ires-eGFP | Gift (M. Nakafuku, Univ. Cincinnati) | N/A |
| Mouse: FVB.B6-Tg(CAG-cat,-EGFP)1Rbns/KrnzJ (CC-eGFP) | The Jackson Laboratory | JAX:024636 |
| Mouse: Tg(Rlbp1-cre/ERT2)1Eml (Rlbp1-CreERT2) | MGI | MGI:7708085 |
| Mouse: B6.Cg-Gt(ROSA)26Sortm14(CAG-tdTomato)Hze/J (RosaAi14) | The Jackson Laboratory | JAX:007914 |
| Mouse: 129-Cdkn1btm1Mlf/J (Cdkn1b flox/flox; p27 flox) | The Jackson Laboratory | JAX:003122 |
| Mouse: H3c6em1Sjes (H3.1-iCOUNT) | The Jackson Laboratory | JAX:037318 |
| Mouse: Hes1tm1(cre/ERT2)Lcm (Hes1:CreERT2) | MGI | MGI:4412375 |
| **Software and algorithms** | | |
| FIJI/ImageJ | NIH | Version 2.9.0 (or later) |
| Imaris | Oxford Instruments | Version 10.2.x |
| GraphPad Prism | GraphPad Software | Version 10.x |
| Zen | Carl Zeiss | Version 2.3 (or later) |
| Trailmaker | Parse Biosciences | Version 1.6.2 |
| Seurat | R Package | 5.4.0 |
| Harmony | R Package | 1.2.4 |
| DoubletFinder | R package | 2.0.6 |
| PIPseeker V pipeline | Fluent BioSciences | V3.3.0 |
| **Other** | | |
| Aurora CS cell sorter | Cytek | S0127 |
| Cell culture insert, 30 mm, hydrophilic PTFE, 0.4 µm | Millicell | PICM0RG50 |
| ECM 830 square wave electroporator | BTX | 45-0052 |
| Platinum disk electrode (7 mm) on stick | Bulldog Bio | CUY700P7L |
| Three-axis coarse mechanical micromanipulator | Narishige | UMM-3C |
| Cytek Aurora CS cell sorter | Cytek | Aurora CS |
| ZEISS Axio Zoom.V16 with Apotome 3 | Carl Zeiss | 495010-0003-000 |
| ZEISS LSM 710 confocal microscope | Carl Zeiss | M60-1-0013 |
| 40 µm cell strainer | Pluriselect | 431004060 |

### Table S2. Electrical stimulation parameters and setup.

| Parameter | Value / description |
|---|---|
| Stimulation timing | 14 DEV (single treatment) |
| Pulse train | Five square-wave pulses, 50 ms each, 250 ms intervals (alternating current) |
| Voltage | 25 V |
| Electroporator | BTX ECM 830 |

| | |
|---|---|
| Electrode type | 7 mm platinum disk electrode (CUY700P7L) |
| Electrode spacing | 6 mm total |
| Calculated field strength | 0.0417 kV/cm |
| Stimulation solution | PBS plus loading dye (see prior method) |
| Retina positioning | Filter center |
| Agarose disk | 0.5% agarose in PBS; 5mm height, 10mm diameter. |

### Table S3. Antibodies and working dilutions.

| Antibody | Dilution | Source | Identifier |
|---|---|---|---|
| Goat α-OTX2 | 1:250 | GeneTex | AF1979 |
| Rabbit α-ASCL1 | 1:250 | Abcam | AB211327 |
| Rabbit α-CABP5 | 1:250 | Sysy | 475002 |
| Rabbit α-cleaved-CASP3 | 1:250 | Biotechne | AF835 |
| Rabbit α-HES1 | 1:250 | Cell SignalingTechnologies | D6P2U |
| Rat α-BrdU | 1:200 | Abcam | AB6326 |
| Alexa Fluor 405 Donkey Anti-Rat IgG (H+L) | 1:500 | Invitrogen | A48268 |
| Alexa Fluor 488 Donkey Anti-Rabbit IgG (H+L) | 1:500 | Invitrogen | A21206 |
| Alexa Fluor 568 Donkey Anti-Rabbit IgG (H+L) | 1:500 | Invitrogen | A10042 |
| Alexa Fluor 594 Donkey Anti-Goat IgG (H+L) | 1:500 | Invitrogen | A11058 |
| Alexa Fluor 647 Donkey Anti-Goat IgG (H+L) | 1:500 | Invitrogen | A21447 |
| Alexa Fluor 647 Donkey Anti-Rabbit IgG (H+L) | 1:500 | Invitrogen | A-31573 |

### Table S4. ScRNA-seq sample metadata and QC Summary.

| Condition & Retina | Mouse & Retina | Mouse age at dissection | Status | Exclusion reason (if excluded) | Cells (post-QC) | Median genes/cell | Median transcripts /cell |
|---|---|---|---|---|---|---|---|
| p27 CKO | 1L | 7 weeks | Excluded | Low RIN | -- | -- | -- |
| p27 CKO | 2R | 7 weeks | Included | | 398 | 5933.5 | 22809 |
| p27CKO | 3L | 9 weeks | Included | | 1677 | 5105 | 16792 |
| p27 CKO | 4R | 9 weeks | Included | | 791 | 3989 | 13507 |
| ESPI | 1R | 7 weeks | Included | | 562 | 5593.5 | 19209 |
| ESPI | 2L | 7 weeks | Excluded | High % mito | -- | -- | -- |
| ESPI | 3R | 9 weeks | Included | | 840 | 4876.5 | 17246 |
| ESPI | 4L | 9 weeks | Included | | 1270 | 5264 | 17569 |

.

### SUPPLEMENTAL FIGURE LEGENDS

**Figure S1. E-stim does not promote expression of downstream neural factors CABP5 & CRX.**
Representative 134.71 μm$^2$ ROIs of 40x whole-mount images showing no overlap between MG-Tom lineage reporter and **(A)** CABP5 or **(B)** CRX expression following E-stim. N=2.

**Figure S2. High density recombination of p27 inactive MG-TOM reporter renders single Müller glial cells indistinguishable for quantification.**
A single 0.54 μm Z-plane, 63X image of p27 inactive MG-Tom retinal tissue after E-stim showing reporter$^+$ Müller glial morphology with BrdU and OTX2 expression.

**Figure S3. Validation of the H3.1-iCOUNT reporter in proliferative embryonic retinal progenitors.**
**(A)** Mouse model and **(B)** timeline schematic for Hes1-CreER$^{T2}$–driven recombination in developing retina to activate the H3.1-iCOUNT, with tamoxifen induction at E16.5 and harvest at E18.5. **(C)** Representative 40x whole-mount images showing robust iCOUNT (H3.1-eGFP) signal in embryonic retina, with absence/unreliable detection of baseline mCherry signal in this tissue. N=3 **(D)** Flow cytometry (FACS) readout from dissociated embryonic retina supporting lack of reliable mCherry detection, motivating downstream analysis based on eGFP intensity. (N=10).

**Figure S4. *p27* WT controls show insufficient iCOUNT eGFP signal for inclusion in iCOUNT$^+$ scRNA-seq sorting.**
**(A)** Müller glia-specific iCOUNT mouse model and **(B)** experimental timeline of iCOUNT$^+$ cell detection in *p27* WT ± E-stim tissues. **(C)** Representative 40x whole mount images demonstrating low iCOUNT (eGFP) detection relative to proliferative labeling, supporting exclusion of *p27* WT conditions from the iCOUNT$^+$ scRNA-seq dataset.

**Figure S5. Microglial clusters identified in the iCOUNT scRNA-seq dataset.**
UMAP visualization showing all clusters including clusters annotated as microglia in the lower right corner (clusters excluded from the main UMAP in Figure 5B), shown for completeness with cluster identities labeled.

**Figure S6. Expression pattern of additional glial, proliferation, and neural markers.**
**(A-C)** Feature plots for Müller glial marker *Glul,* and for *Pax6* and *Vsx2* genes, which are expressed in adult Müller glia and retinal progenitor cells during development. **(D-F)** Feature plots for the NFI genes *Nfia, Nfib*, and *Nfix.* **(G, H)** Feature plots for rod photoreceptor-associated genes (*Pde6g, Nr2e3*).

**Figure S7. Assessment of apoptosis after electrical stimulation in *p27* CKO iCOUNT retinas.**
**(A)** Mouse model and **(B)** experimental timeline matching Figure 7 for *p27* CKO H3.1-iCOUNT retinas cultured *ex vivo* ± E-stim with extended BrdU delivery post-E-stim and fixation at 20 DEV. **(C)**

Representative 40x images of whole mount retina immunostaining for cleaved CASP3 in iCOUNT$^+$ Müller glia shows minimal evidence of apoptosis in unstimulated *p27* CKO and *p27* CKO + E-stim retinas.
