## Supplementary figures and images for "Electrical stimulation combined with p27Kip1 inactivation drives proliferative neurogenic reprogramming of Mueller glia in the adult mouse retina"

### Figure S1

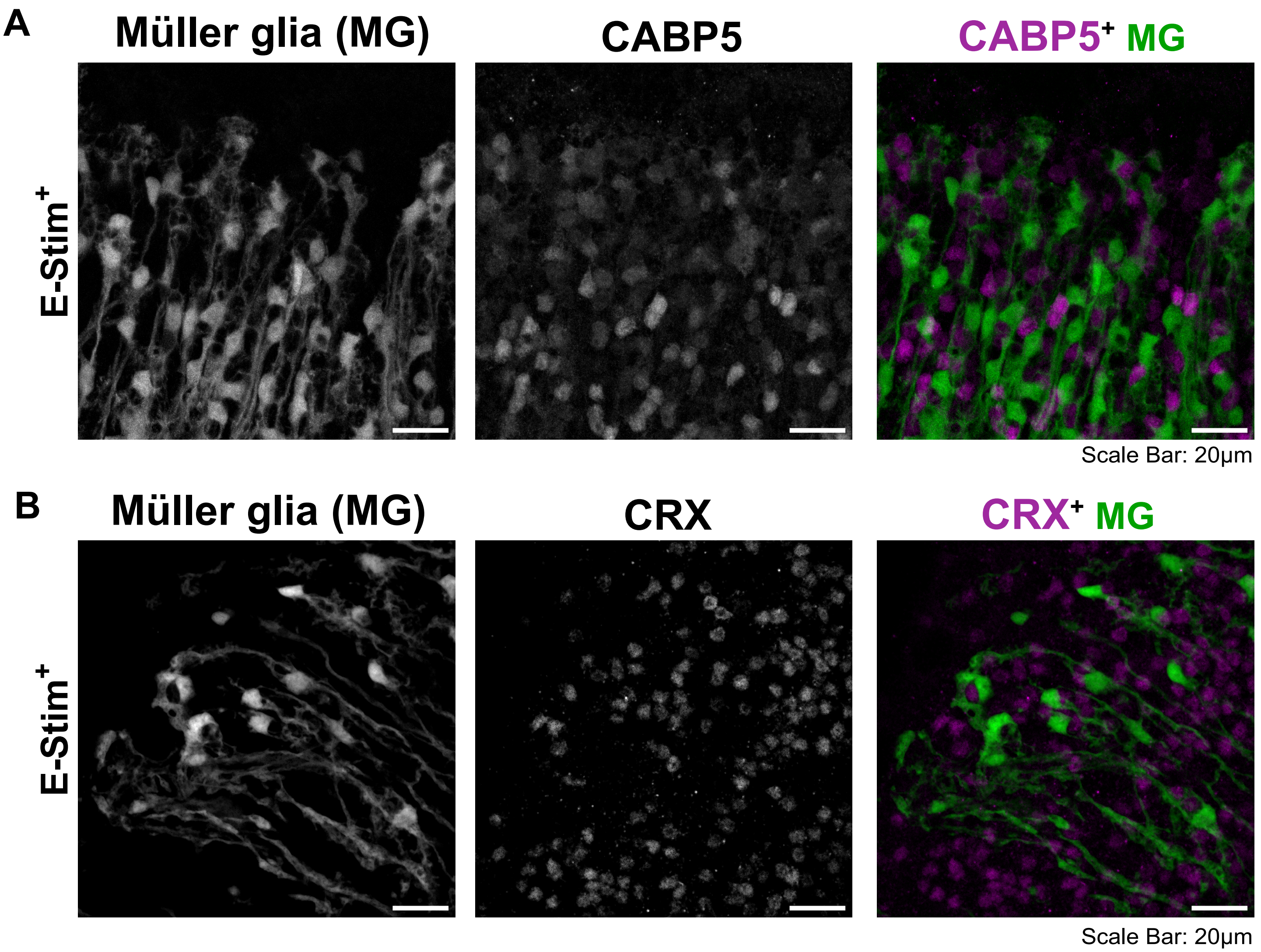

### Figure S2

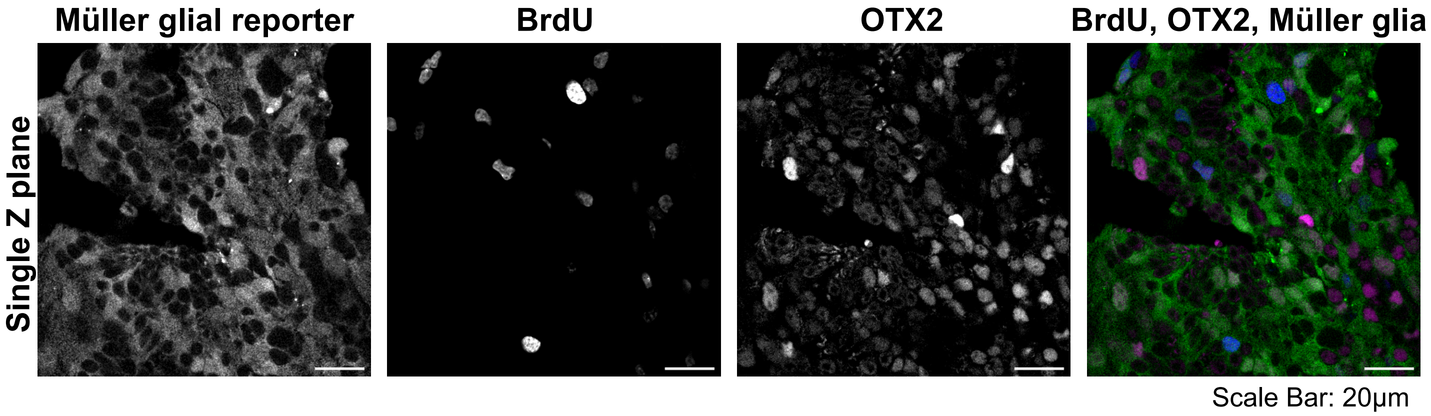

### Figure S4

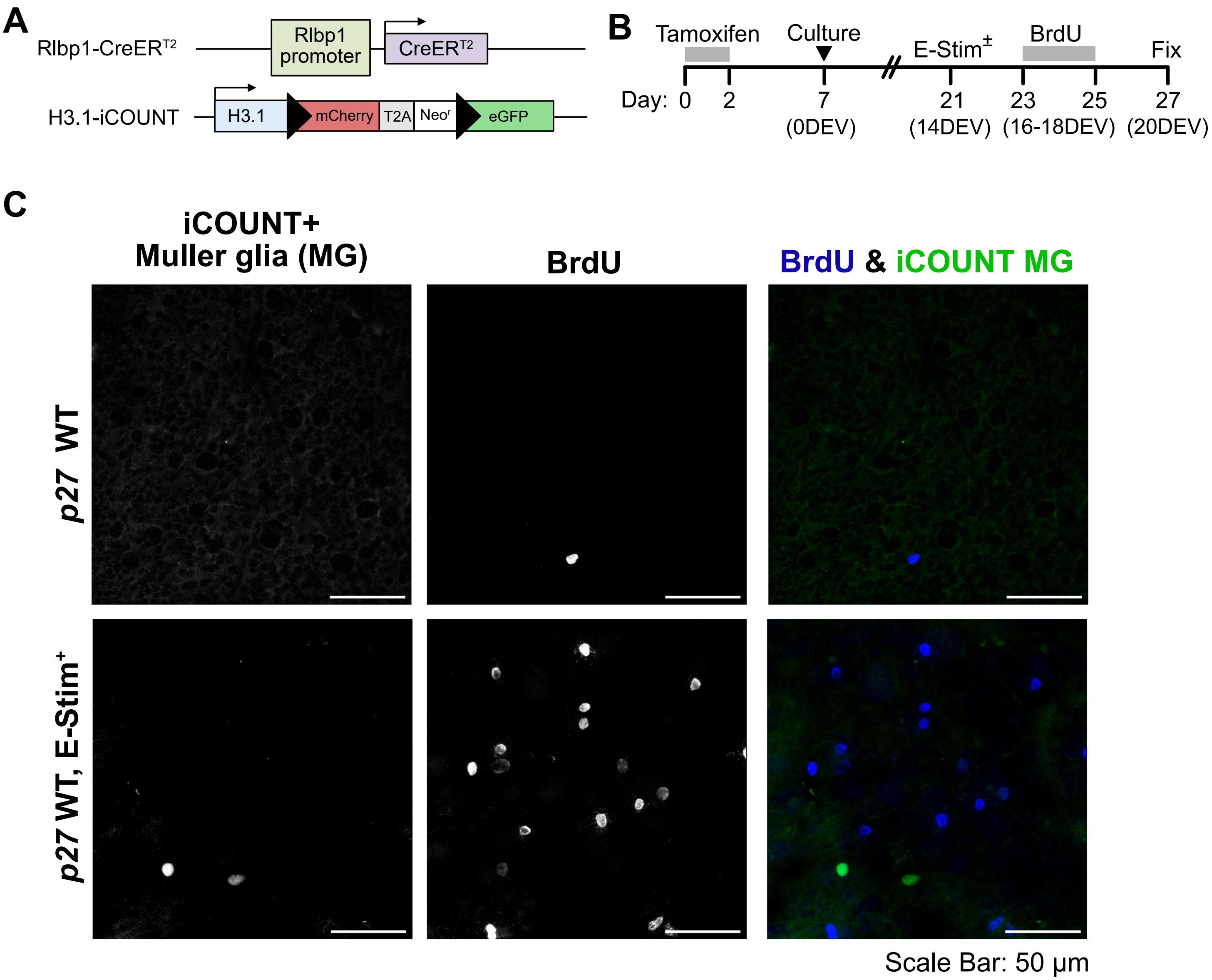

### Figure S5

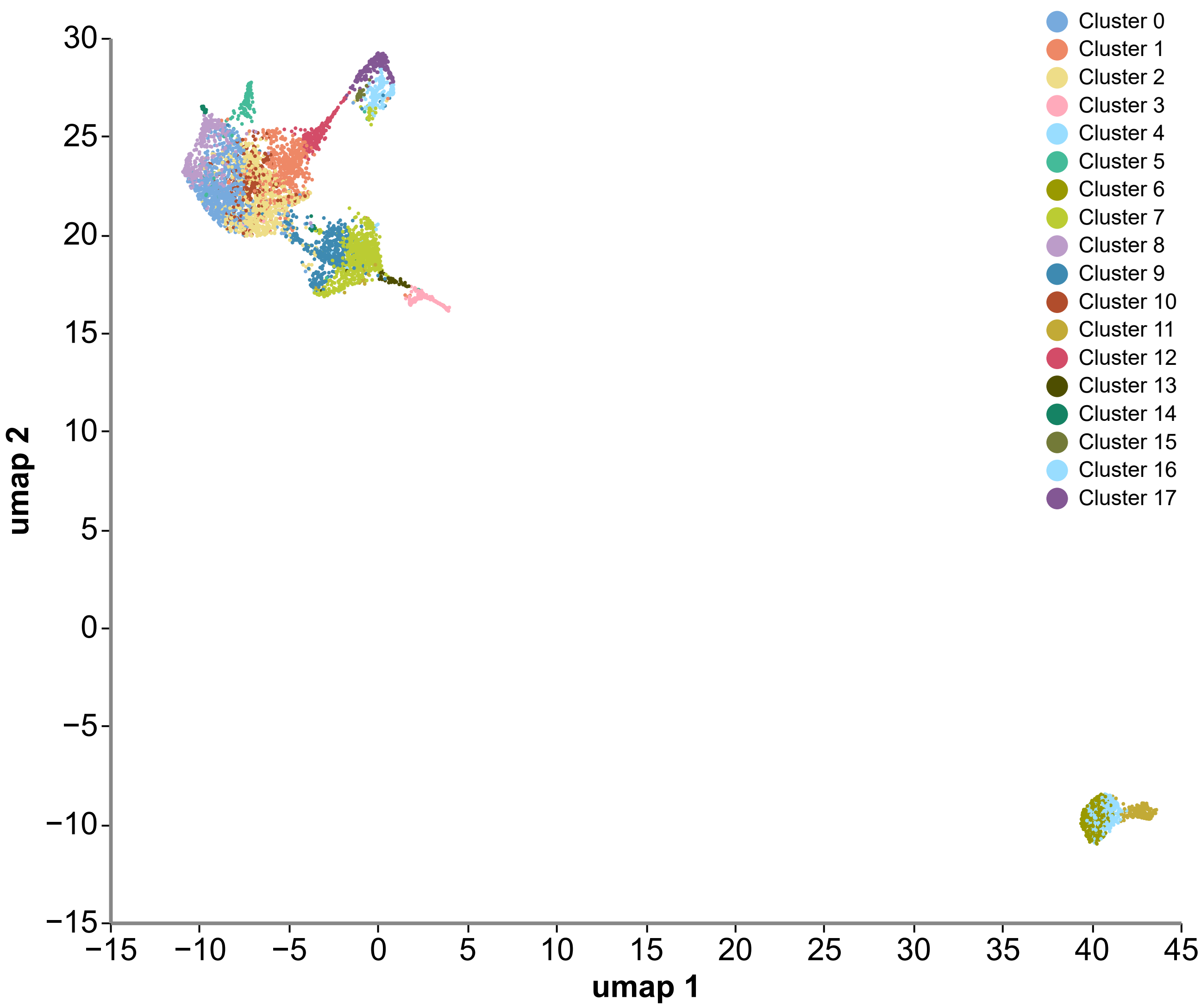

### Figure S6

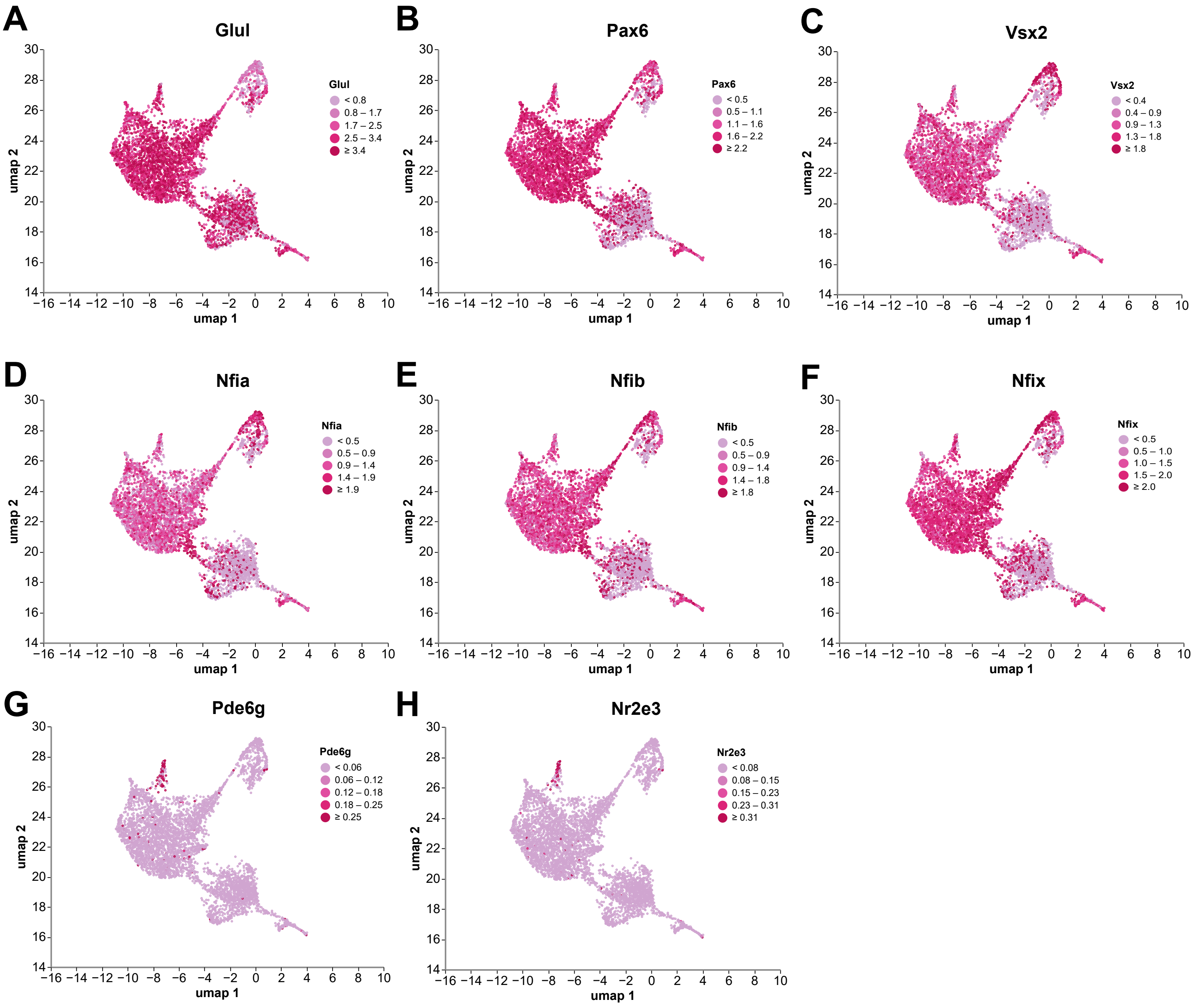

### Figure S7

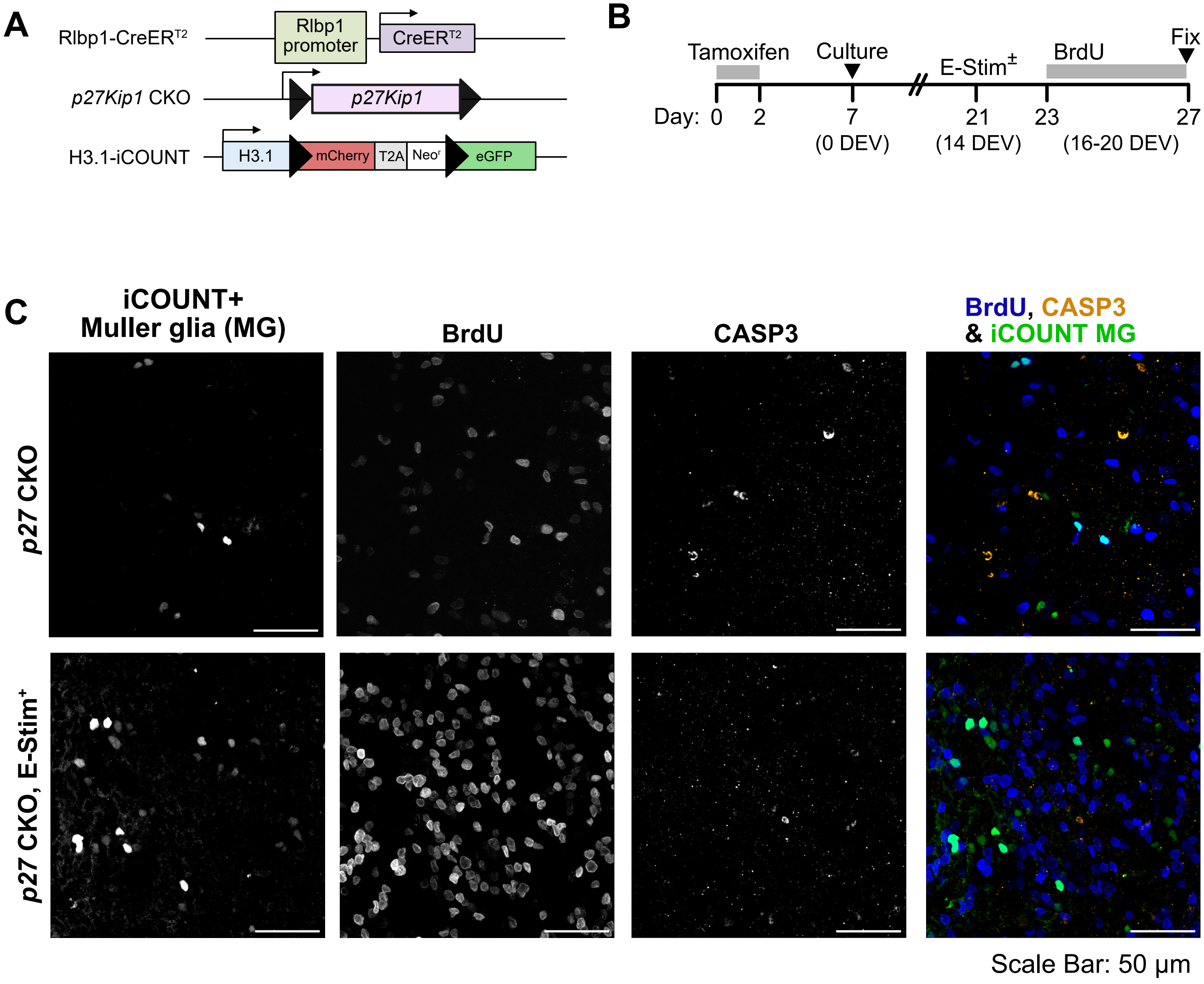
