## Supplementary material for "Electrical stimulation combined with p27Kip1 inactivation drives proliferative neurogenic reprogramming of Mueller glia in the adult mouse retina": Figure S3

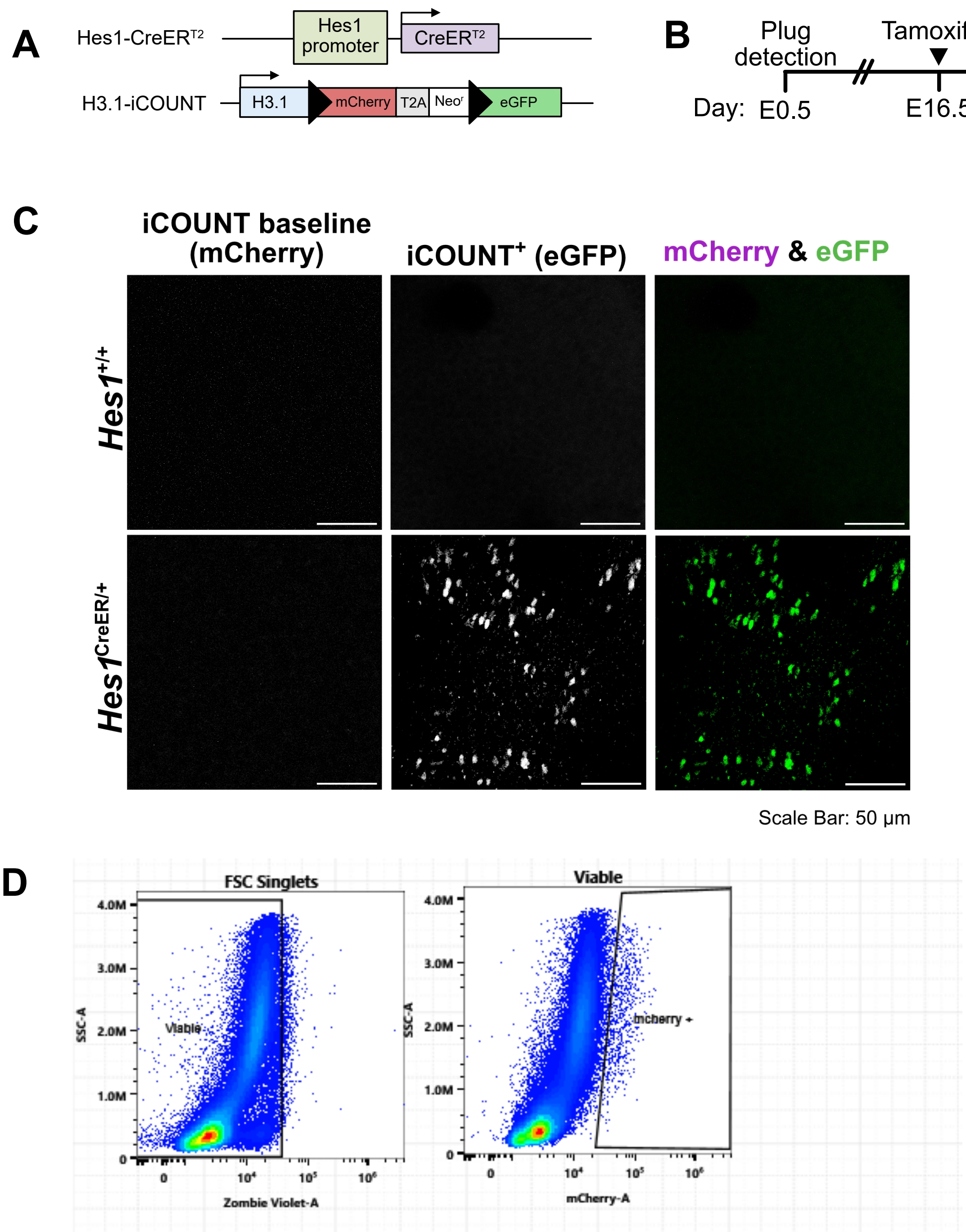


Population Hierarchy
Levine_Stone_072225-Group_001-C1

| Population | % Parent | Count |
|---|---|---|
| All Events | 100.00 | 271,335 |
| Scatter | 66.14 | 179,472 |
| SSC Singlets | 97.77 | 175,462 |
| FSC Singlets | 98.92 | 173,562 |
| Viable | 98.48 | 170,924 |
| mcherry + | 0.96 | 1,635 |
